# Knowledge-primed neural networks enable biologically interpretable deep learning on single-cell sequencing data

**DOI:** 10.1101/794503

**Authors:** Nikolaus Fortelny, Christoph Bock

## Abstract

Deep learning has emerged as a powerful methodology for predicting a variety of complex biological phenomena. However, its utility for biological discovery has so far been limited, given that generic deep neural networks provide little insight into the biological mechanisms that underlie a successful prediction. Here we demonstrate deep learning on biological networks, where every node has a molecular equivalent (such as a protein or gene) and every edge has a mechanistic interpretation (e.g., a regulatory interaction along a signaling pathway).

With knowledge-primed neural networks (KPNNs), we exploit the ability of deep learning algorithms to assign meaningful weights to multi-layered networks for interpretable deep learning. We introduce three methodological advances in the learning algorithm that enhance interpretability of the learnt KPNNs: Stabilizing node weights in the presence of redundancy, enhancing the quantitative interpretability of node weights, and controlling for the uneven connectivity inherent to biological networks. We demonstrate the power of our approach on two single-cell RNA-seq datasets, predicting T cell receptor stimulation in a standardized *in vitro* model and inferring cell type in Human Cell Atlas reference data comprising 483,084 immune cells.

In summary, we introduce KPNNs as a method that combines the predictive power of deep learning with the interpretability of biological networks. While demonstrated here on single-cell sequencing data, this method is broadly relevant to other research areas where prior domain knowledge can be represented as networks.

## INTRODUCTION

Deep learning using artificial neural networks (ANNs) has achieved unprecedented prediction performance for complex tasks in multiple fields, including image recognition^1–3^, speech recognition^4,5^, natural language processing and machine translation^6–10^, board and computer games^11–13^, and autonomous driving^14,15^. There is tremendous potential for deep learning in biology and medicine^16–19^, as illustrated by initial applications in medical image classification^20^, brain image segmentation^21^, epigenome prediction^22^, DNA/RNA binding analysis^23^, and RNA splicing inference^24,25^. Most recently, deep learning has shown promising results in the analysis of single-cell RNA sequencing (RNA-seq) datasets, facilitated by the large number of assayed cells^26–30^.

For specialized tasks, ANNs have achieved performance on par with, or superior to, that of human experts. However, the trained ANNs typically lack interpretability, i.e., the ability to provide human-understandable, high-level explanations of how they transform inputs (prediction attributes) into outputs (predicted class values). Indeed, ANNs are structurally very different from biological networks, and no molecular equivalent exists for the internal nodes and edges of an ANN trained on a biological dataset. As the result, there is no obvious way to biologically interpret trained ANNs. This lack of interpretability in deep learning is a major limitation to the wider application of this powerful technology in biology and medicine – not only because it reduces trust and confidence in using deep learning predictions for high-stakes applications such as clinical diagnostics^17,18^, but also because it misses important opportunities for data-driven biological discovery using deep learning.

Pioneering research aimed at making deep learning models interpretable and informative for biological applications has focused on the *ex post* analysis of trained ANNs, for example by identifying inputs that result in specific predictions^23,31,32^ or analyzing compressed layers in autoencoders^33^. As an alternative and complementary strategy, it may be possible to engineer deep learning methods for built-in interpretability, for example by incorporating domain knowledge into the ANNs used for deep learning. Initial applications of this approach have embedded protein interactions into autoencoders^34^ and derived ANNs from gene annotations^35^.

Embedding of biological knowledge into ANNs creates an opportunity for systems biological modeling by deep learning. Biological processes are controlled by the interplay of biological entities (such as genes or proteins) that cooperate to maintain complex biological phenotypes. This cellular logic is captured by biological networks^36,37^, for example describing the signaling pathways and transcription-regulatory mechanisms that control homeostatic cell state and response to stimuli. Cells can thus be conceived as living “information processing units”^38,39^ that perform calculations on biological networks in order to regulate cell state. Computational models that predict cell state from molecular data approximate these “biological calculations”. In other words, when we train ANNs to predict cellular phenotypes based on high-throughput genomics data, we optimize artificial networks in the computer to approximate calculations that cells perform through biological networks *in vivo*.

To overcome lack of interpretability as a key limitation of deep learning in biology and medicine, here we explore the concept – and demonstrate the feasibility – of deep learning on application-specific biological networks rather than on generic ANNs. By applying deep learning directly to biological networks (such as signaling pathways and gene-regulatory networks), this approach coerces deep learning algorithms to stick closely to the regulatory mechanisms that are relevant in cells, thereby making the learned models interpretable.

We developed knowledge-primed neural networks (KPNNs) as a framework for interpretable deep learning on biological networks (applied here to signaling pathways and gene-regulatory networks), and we introduce three modifications to generic deep learning that enhance interpretability: (i) Repeated network training with random deletion of hidden nodes (a technique known as dropout^40^), which yields robust results in the presence of network redundancy; (ii) dropout on input data in order to enhance quantitative interpretability of node weights; (iii) training on control inputs to normalize for the uneven connectivity of biological networks. To validate our method for interpretable deep learning on biological networks, we applied KPNNs to single-cell RNA-seq data for T cell receptor stimulation^41^ and to a reference catalog of immune cells from the Human Cell Atlas^42^.

## RESULTS

### KPNNs enable deep learning on biological networks

Deep learning using artificial neural networks (ANNs) seeks to “learn” (i.e., approximate in a generalizable way) the complex relationship between a set of prediction attributes (such as single-cell transcriptomes) and the corresponding class values (such as cell type or cell signaling state). The learning process typically starts from a generic, fully connected, feedforward ANN – which is a directed acyclic graph organized in layers such that each node receives input from all nodes in the previous layer and sends its output to all nodes in the next layer. During the learning process, edge weights are randomly initiated and iteratively updated based on training data, seeking to improve the accuracy with which the ANN transforms the inputs into corresponding class values. With enough training data, large multi-layer ANNs can learn highly complex relationships between prediction attributes and class values, without requiring any domain knowledge. However, the resulting trained ANNs lack interpretability, given that their nodes, edges, and weights do not correspond to meaningful domain concepts. They are also inherently instable, such that very distinct networks can achieve similar performance.

To overcome the lack of interpretability of deep learning on ANNs, we sought to embed relevant biological knowledge into neural networks that are trained by deep learning (**Fig. 1**). We replaced the fully connected networks in ANNs – the substrate of deep learning – by networks derived from biological knowledge, thereby creating “knowledge-primed neural networks” (KPNNs). In KPNNs, each node corresponds to either a protein or a gene, and each edge corresponds to a potential regulatory relationship that has been experimentally observed and annotated in public databases. We constructed KPNNs to reflect the typical flow of information in cells, where signals are transduced from receptors via signaling proteins to transcription factors, which in turn induce changes in gene expression. For deep learning on KPNNs, the expression levels of the regulated genes are modeled as input nodes and are experimentally measured. In contrast, signaling proteins and transcription factors are modeled as hidden nodes, and their activation states are inferred during the learning process. Finally, receptors are connected to output nodes for complex multi-class prediction (e.g. classifying cell types) or themselves represent the output node (e.g., predicting receptor-stimulated vs unstimulated cells).

**Figure 1.**
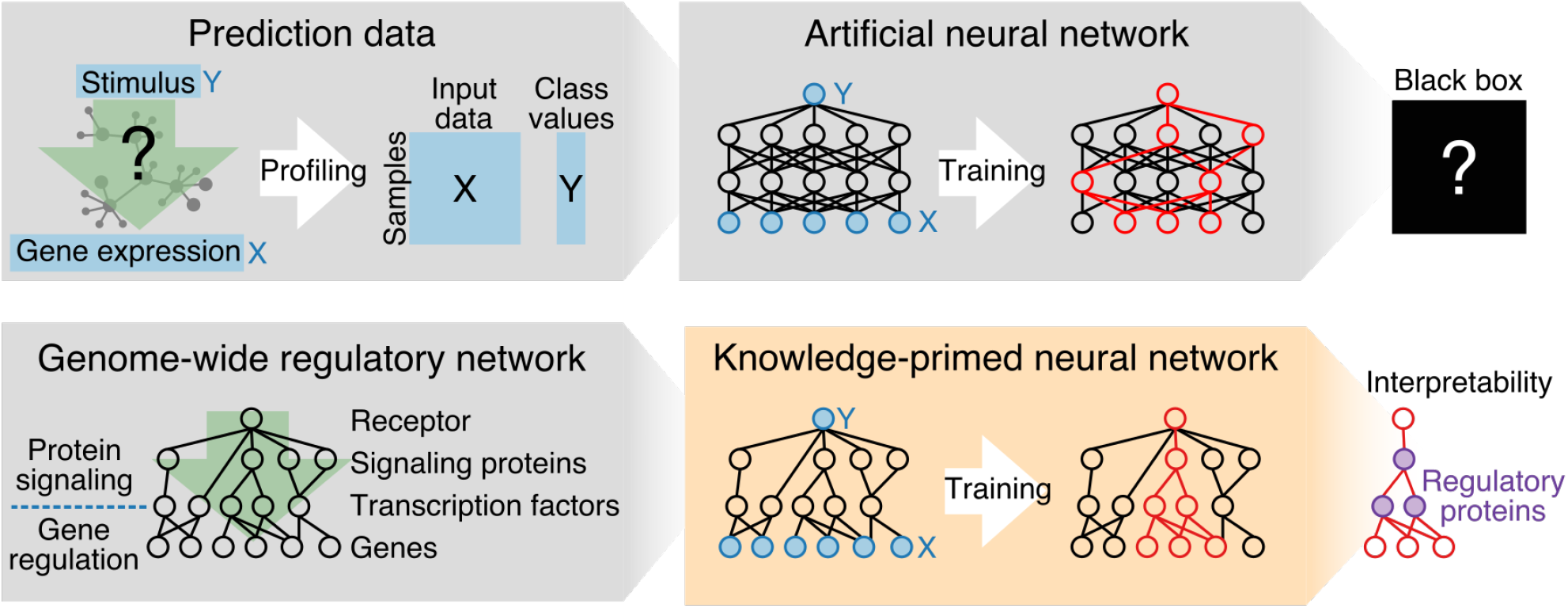
Interpretable deep learning with KPNNs. Deep learning provides a powerful method for predicting cell states from gene expression data. However, prediction using artificial neural networks (ANNs, top row) is a “black box” method that provides little insight into the biology that underlies a successful prediction – for two reasons: (i) Nodes and edges in an ANN have no biological equivalent and therefore lack interpretability; (ii) ANNs are inherently instable, and very different networks can achieve similar performance. Knowledge-primed neural networks (KPNNs, bottom row) enable interpretable deep learning on biological networks by exploiting structural analogies between biological networks (such as the signaling pathways and gene-regulatory networks that regulate cell state) and the feed-forward neural networks used for deep learning. In KPNNs, each hidden node corresponds to a regulator protein, and each edge corresponds to a potential regulatory relationship that has been reported in public databases. Exploiting the sparsity of KPNNs and the power of deep learning to assign meaningful weights across multi-layer networks, it is possible to train KPNNs in such a way that the learnt weights are biologically interpretable as robust estimates of regulatory protein activity.

KPNNs can be trained in much the same way as ANNs (see Methods for details), for example based on training data that comprise gene expression profiles for cells of different type (when classifying cell types) or exposed to different conditions (when predicting receptor-stimulated vs unstimulated cells). After successful completion of the learning process, the fitted edge weights estimate the relevance of the corresponding potential regulatory relationships in the biological system under investigation. Furthermore, we can derive node weights from the edge weights, in order to identify the most relevant signaling proteins and regulatory factors. Conceptually, interpretable deep learning with KPNNs is based on two assumptions: (i) most of the potential regulatory relationships that are important in a biological system of interest have been observed previously in other biological contexts and are annotated in public databases of signaling pathways and gene-regulatory networks; and (ii) deep learning makes it possible to assign meaningful weights across multiple hidden layers in KPNNs, thereby providing useful estimates of regulatory protein activity (which is often difficult to measure experimentally).

We applied and evaluated the KPNN method in two initial biological applications: Dissecting cellular response to T cell receptor (TCR) stimulation based on single-cell RNA-seq data (1,735 single-cell transcriptomes)^41^ and inferring cell type in a Human Cell Atlas (HCA) reference dataset comprising 483,084 immune cells^42^ (**Supplemental Fig. 1**). To construct the TCR KPNN, we connected the TCR (output node) to single-cell gene expression profiles (inputs) via shortest paths through a network of gene-regulatory and protein signaling interactions (hidden nodes). The resulting network was then reversed and trained to predict TCR stimulation from gene expression: Gene expression (input nodes) provides the input for the activity state of transcription factors (hidden nodes), whose outputs are used by signaling proteins (hidden nodes) to predict TCR stimulation (output node). For the HCA KPNN, we combined and integrated various signaling pathways into a single network. Single-cell gene expression profiles (input nodes) were connected to cell surface receptors (hidden nodes) via a network of gene-regulatory and protein signaling interactions (hidden nodes). Cell surface receptors were then connected to three nodes representing B cells, T cells, and monocytes (output nodes).

In both applications, deep learning on KPNNs provided high prediction accuracies comparable to that of deep learning using ANNs (**Supplemental Fig. 2**). The TCR KPNN predicted TCR stimulation with a median receiver operating characteristic (ROC) area under curve (AUC) value of 0.984 (inter-quartile range: 0.979 to 0.987), while ANNs with the same number of nodes (and many more edges) achieved a median ROC AUC value of 0.948 to 0.985 depending on the number of layers (interquartile range: 0.936 to 0.988 across all analyses; 0.938 to 0.989 for the best-performing number of layers). Similarly, the HCA KPNN predicted immune cell types in the HCA dataset with a median ROC AUC value of 0.994 (inter-quartile range: 0.992 to 0.995), while ANNs with the same number of nodes (and many more edges) achieved a median ROC AUC value of 0.5 to 0.995 depending on the number of layers (interquartile range: 0.5 to 0.992 across all analyses; 0.993 to 0.996 for the best-performing number of layers).

In summary, we demonstrated that deep learning on KPNNs is feasible and can achieve comparable accuracies to deep learning on ANNs. This result holds the promise that deep learning on restricted networks with a direct molecular equivalent for nodes and edges may yield network models that are accurate and interpretable.

### The biology-based structure of KPNNs fosters interpretability

The interpretability of KPNN relies in part on the specific structure of biological networks, which are much sparser and have lower complexity than ANNs with the same number of nodes. For example, the best-performing ANNs had 151-fold (TCR) and 210-fold (HCA) more edges than the corresponding KPNN (**Supplemental Fig. 3**). To investigate the structural differences of KPNNs and ANNs, we systematically compared network properties of our two KPNNs (TCA, HCA) to fully connected ANNs with the same number of nodes (fANNs), and to sparse ANNs with the same number of edges as the KPNNs (sANNs) obtained by a layer-wise, random deletion of edges from the fully connected ANNs (see Methods for details).

Beyond different edge densities, the most characteristic structural difference between KPNNs and ANNs was the prevalence of deep connections (“shortcuts”) in KPNNs, contrasting with the strictly layered network architecture of ANNs (**Fig. 2a, Supplemental Fig. 4a and 5a**). Because each node in the fully connected ANNs is connected to all nodes of the previous and following layers, the distance from a given node to any node of the input layer is equal and depends only on the node’s layer. In contrast, distances in KPNN differ widely due to the presence of shortcuts that connect certain parts of the network much more directly with the input layer.

**Figure 2.**
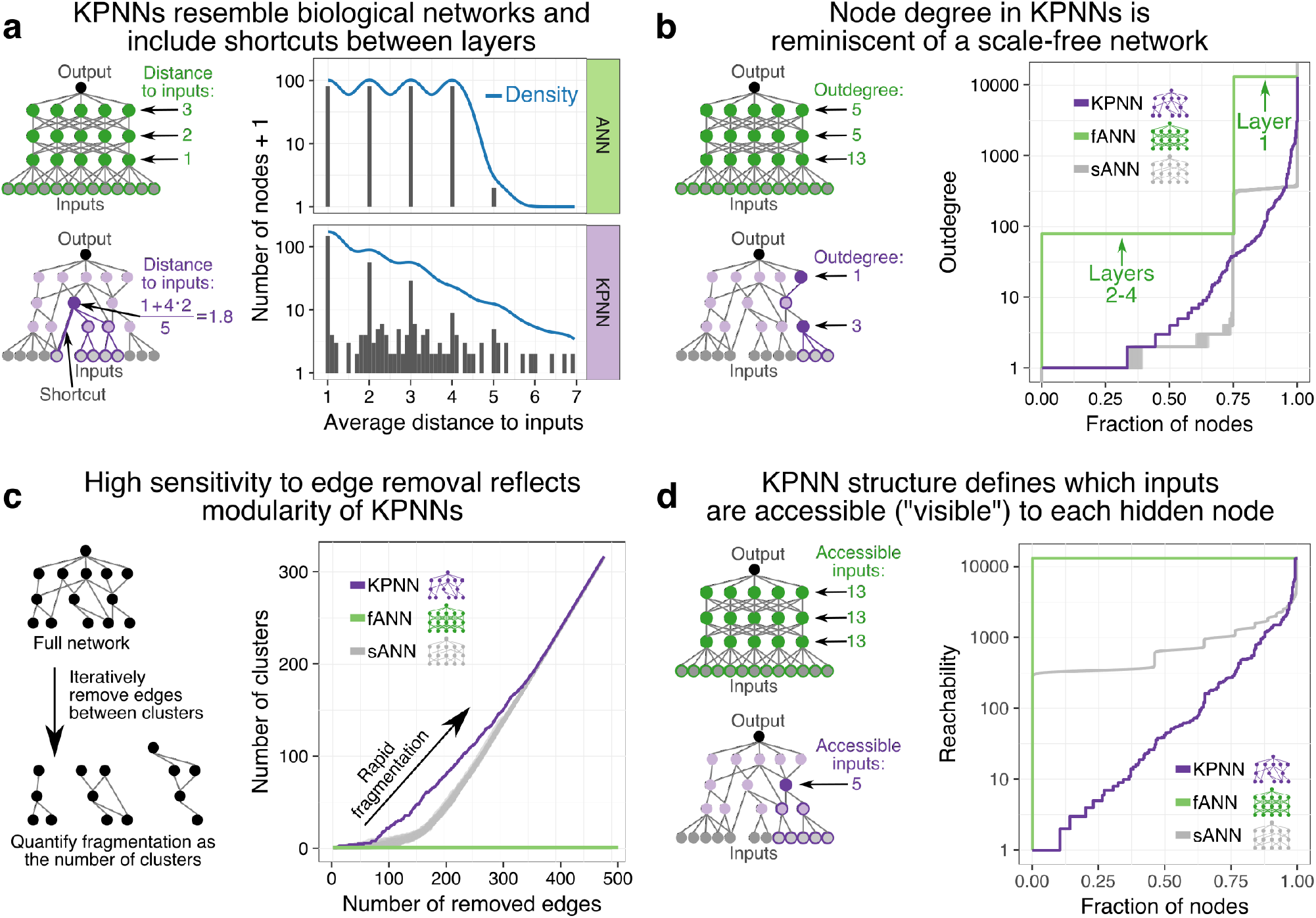
Comparative network analysis of KPNNs and ANNs. The TCR KPNN is compared to a fully connected ANN (fANN) with the same number of nodes and the same median depth as the KPNN (4 hidden layers), and to sparse ANNs (sANNs) where edges were randomly removed to match the edge number of the KPNN (results are shown for 50 random sANNs). **(a)** Average distance to the input nodes for all hidden nodes in the fANN (top) and KPNN (bottom). **(b)** Cumulative distribution of the outdegree of hidden nodes in the KPNN, the fANN, and the sANNs. **(c)** Assessment of network sensitivity to fragmentation upon removal of important edges in the KPNN, the fANN, and the sANNs. Edges were iteratively removed based on importance measured by their network betweenness value. **(d)** Cumulative distribution of reachability, which measures the number of input nodes each hidden node can connect to, shown for the KPNN, the fANN, and the sANNs.

KPNNs are further characterized by their patterns of connectedness, as illustrated by the distribution of node outdegrees (the outdegree of a node measures the number of edges that leave the node). The outdegrees in the KPNNs resembled an exponential distribution; in contrast, nodes in the ANNs are all connected to the same (constant) number of other nodes (**Fig. 2b, Supplemental Fig. 4b,c and 5b,c**). The approximately scale-free nature of KPNNs is consistent with the well-established property of biological networks having a few highly connected “hub” nodes that link to many nodes with lower connectivity^39^. As a result of these connectivity patterns, KPNNs also showed a higher degree of modularity than ANNs, demonstrated by their substantially higher sensitivity to fragmentation upon edge removal compared to fully connected ANNs, and even to sparse ANNs (**Fig. 2c, Supplemental Fig. 4d and 5d**).

The sparseness and modularity of KPNNs restricts the information (i.e., input nodes) that are accessible to each hidden node through direct or indirect connections. This is in contrast to fully connected ANNs, where each hidden node can access all input nodes. To quantify the amount of information available to each hidden node, we calculated the network metric “reachability”, i.e. the number of input nodes that a given hidden node can access through direct or indirect connections. Strikingly, most hidden nodes in KPNNs were connected only to a small fraction of input nodes. The restricted access to input information was less pronounced in edge-matched sparse ANNs and absent from fully connected ANNs (**Fig. 2d, Supplemental Fig. 4e and 5e**).

In summary, the network structure of KPNNs strongly deviates from that of generic ANNs. It rather reflects the specific characteristics of biological networks, which constrain the flow of information through the network by a sparse, modular architecture that includes many shortcuts. These characteristic network properties were qualitatively and quantitatively similar between the TCR KPNN and the HCA KPNN. They can be expected to benefit the biological interpretability of KPNNs because they dramatically reduce the free parameter space and force the trained KPNNs to approximate biologically relevant regulatory mechanisms. Reassuringly, these restrictions did not come at the expense of low prediction accuracies (**Supplemental Fig. 2**), which reinforces the hypothesis that KPNNs indeed capture the underlying biology well enough to support interpretability.

### An optimized learning method renders KPNNs interpretable

Interpretable deep learning using KPNNs relies on two key components – the biological network structure, which is based on a broad range of possible regulatory relationships identified by prior research and obtained from public databases; and a deep learning process that determines which elements of the KPNN are most relevant in the biological system under investigation. Specifically, the TCR KPNN and the HCA KPNN were both derived from large databases comprising gene-regulatory interactions and protein signaling pathways (independent of the concrete training data), and the resulting KPNNs capture a wide range of biological processes in one network. Training of the KPNNs using single-cell RNA-seq data iteratively updates edge weights in the networks until they reflect the relationship of input data to output class values in the biological system under investigation. Finally, we calculate hidden node weights and interpret them as estimates of regulatory protein activity of the corresponding signaling protein or regulatory factor.

While generic deep learning algorithms achieve high prediction accuracies for KPNNs (**Supplemental Fig. 2**), we identified three major obstacles to interpretable deep learning on KPNNs: (i) insufficient reproducibility, especially in the presence of network redundancy; (ii) limited quantitative interpretability, where node weights fail to adequately reflect differences in the predictive power of different nodes; and (iii) uneven connectivity inherent to biological networks, which affects node weights independent of training data. To address these issues, we developed an optimized learning method for KPNNs that enhances their interpretability.

First, biological networks are characterized by widespread redundancy, for example when one protein regulates another protein via two separate signaling pathways. As the result, widely different trained models for the same network may achieve similar prediction performance, and generic deep learning algorithms can result in highly divergent weights^33^. Such model variability is not a major issue for conventional applications of deep learning, where the focus is on achieving high prediction performance, while little attention is given to individual weights. However, it has the potential to undermine interpretability of KPNNs, because identical networks trained on identical data can result in highly variable weights and thereby yield inconsistent interpretations^43,44^.

To illustrate this problem and to demonstrate our solution, we simulated and analyzed a series of simple gene-regulatory networks with matched training data (see Methods for details). We first engineered a network without any node redundancy, performed model fitting, and then calculated node weights. In this redundancy-free network, node weights accurately reflected the information flow that we embedded into the network (**Fig. 3a**). Next, when we introduced redundancy into the network and repeatedly trained the model, the standard learning algorithm (see Methods for details) produced node weights that were highly variable (**Fig. 3b**). Strikingly, node weights of redundant nodes were inversely correlated (R = −0.79) across network replicates. Network replicates thus led to inconsistent and therefore uninterpretable results using the standard learning algorithm.

**Figure 3.**
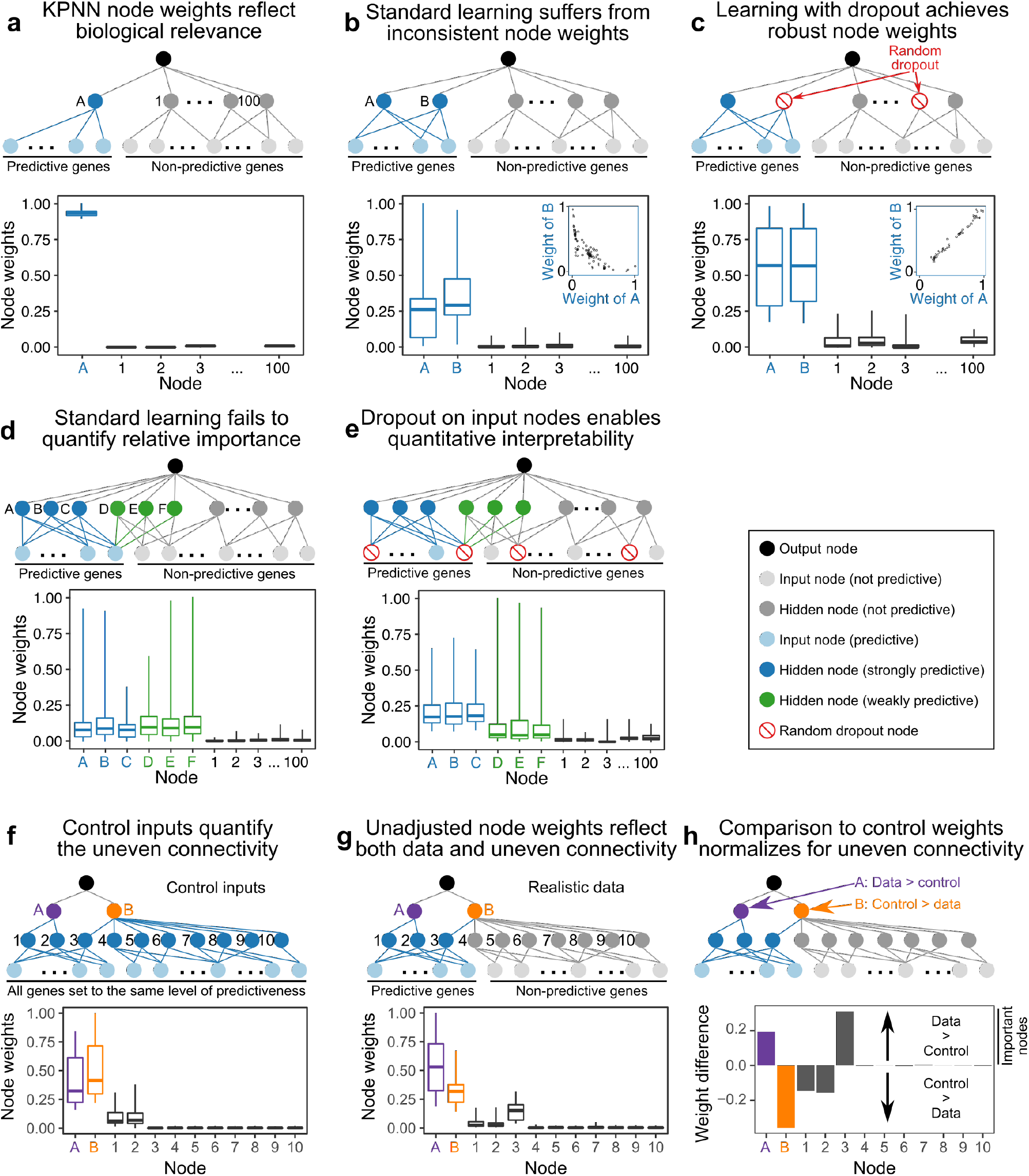
Optimized learning methodology for KPNNs. **(a)** Node weights reflect node importance in a simple network. (Top) Simulated network with one hidden node (node A) that is connected to predictive input nodes. (Bottom) Node weights distinguish predictive from non-predictive nodes after network training. **(b)** High variability of node weights for two redundant nodes based on generic deep learning. (Top) Network with two hidden nodes (A and B) connected to predictive input nodes. (Bottom) Node weights distinguish predictive from non-predictive nodes, but there is a negative correlation of node weights for the two redundant nodes (inset). **(c)** Dropout reduces variability and increases robustness of node weights. (Top) The same network as in (b), trained with dropout on hidden nodes. While two nodes with dropout are indicated as examples, different dropout nodes are randomly selected for each sample and at each training iteration. (Bottom) Learning with dropout results in robust and highly correlated (inset) weights for the two redundant nodes. **(d)** Node weights of weakly and strongly predictive nodes using generic deep learning. (Top) Network with three strongly predictive hidden nodes (A-C, connected to multiple predictive input nodes) and three weakly predictive hidden nodes (D-F, connected to one predictive input node). (Bottom) Node weights do not separate highly predictive from weakly predictive nodes using generic deep learning. **(e)** Learning with input node dropout distinguishes between highly predictive and weakly predictive hidden nodes. (Top) The same network as in (d), trained with dropout on input nodes. (Bottom) Node weights separate highly predictive from weakly predictive nodes when training with input node dropout. **(f)** Control inputs quantify the uneven connectivity of biological networks. (Top) Network with two layers of hidden nodes (A and B; 1 to 10) and input nodes that are all predictive of the output. (Bottom) Node weights trained on control inputs reflect the uneven connectivity of the simulated network. **(g)** Node weights obtained by training on actual data reflect both the data and the uneven connectivity. (Top) The same network as in (f), but with only a subset of input nodes being predictive. (Bottom) Node weights for the network trained on actual data. **(h)** Comparison of node weights for actual data and for control inputs enables normalization for uneven network connectivity. (Top) The same network as in (g), with annotation of the effect of input data and network structure on the importance of nodes A and B. (Bottom) Differential node weights for actual data versus control inputs.

To address this issue and to obtain robust, interpretable node weights despite network redundancy, we incorporated random dropout of hidden nodes into our learning procedure. When networks are trained with dropout, a given percentage of nodes is randomly selected and “dropped” (i.e., set to zero) for each sample and training step (see Methods for details). Deep learning with dropout was originally proposed as a strategy to improve generalizability of ANNs^40^. We found that random dropout of hidden nodes dramatically improved consistency of node weights across network replicates, resulting in an almost perfect correlation of weights for redundant nodes (R = 0.99, **Fig. 3c**) and an improved overall correlation of node weights between network replicates (**Supplemental Fig. 6**). These results demonstrate that dropout of hidden nodes forces deep learning to spread weights across the network instead of depending on individual nodes, which improves robustness.

Second, we found that node weights obtained by standard learning algorithms did not adequately reflect differences in the predictive power of different nodes. We illustrate this issue using a simulated network comprising three strongly predictive hidden nodes (these nodes were connected to several predictive input nodes) and three weakly predictive hidden nodes (connected to just one predictive input node). Using standard learning, this difference was not reflected in the trained node weights (**Fig. 3d**). To address this issue, we introduced dropout on input nodes into the learning procedure (in addition to dropout on hidden nodes), thereby forcing the model to spread weights across input nodes. As a result, we indeed observed much-improved quantitative interpretability, with a clear difference in node weights between nodes with different predictive power (**Fig. 3e**).

The third obstacle for interpretable deep learning on KPNNs lies in the uneven connectivity inherent to biological networks, where some proteins are much more strongly connected than others^39,45^. The network structure affects the trained weights independently of input data, for example by preferentially assigning high weights to central, well-connected nodes. To illustrate the effect of uneven connectivity on node weights, we simulated a network with 12 hidden nodes organized into a top layer (nodes A and B) and a bottom layer (nodes 1 to 10). We constructed this network with uneven connectivity, such that one of the top-level nodes is connected to more nodes than the other (A: 2 nodes, B: 8 nodes). Next, we trained this network with control inputs that were simulated such that all input nodes are informative (predictive of the output). Networks trained on these controls reflected the embedded connectivity patterns: The more strongly connected node B received a higher node weight than the less connected node A; and more central nodes in the top layer (node A and B) received higher node weights than the nodes in the bottom layer (nodes 1 to 10) (**Fig. 3f**).

To normalize for uneven connectivity in KPNNs, we compared models that were trained on realistic data (resulting in node weights that reflect both the simulated biological signal and the effect of uneven connectivity, **Fig. 3g**) to models trained on control inputs (resulting in node weights that solely reflect network connectivity, **Fig. 3f**). By comparing node weights between these two analyses, we can quantify the degree to which a given node is more or less important than expected based on its network connectivity. The resulting “differential node weights” thus provide a normalized measure of node importance in the training data that is adjusted for the effect that the KPNN structure would otherwise have on trained node weights. Differential node weights clearly distinguished the respective data-driven and connectivity-driven weights of nodes A (with 2 of 2 downstream nodes predictive) and B (with only 1 of 5 downstream nodes predictive) in our simulation (**Fig. 3h**).

In summary, we introduced three methodological improvements for interpretable deep learning on KPNNs: hidden node dropout enhances robustness of node weights in the presence of network redundancy; input node dropout distinguishes between strongly predictive and weakly predictive nodes; and the calculation of node weights based on control inputs allows us to normalize for the effect of uneven connectivity in biological networks. In combination with the sparse structure of KPNNs and the fact that every node and edge in the KPNNs has a molecular equivalent, this approach creates a powerful methodology for interpretable deep learning.

### KPNNs establish interpretable prediction models of T cell receptor stimulation

As the first biological test case for interpretable deep learning using KPNNs, we chose our recently published single-cell RNA-seq dataset measuring the cellular response to T cell receptor (TCR) stimulation in a standardized *in vitro* model^41^. The TCR signaling pathway, which orchestrates the transcriptional response to antigen detection in T cells, is well-suited for evaluating the practical utility of our method, given the pathway’s complexity and its well-characterized nature.

We observed excellent performance when predicting from transcriptome profiles whether or not single cells were TCR stimulated, using both the standard learning method applied to KPNNs (**Supplemental Fig. 2a**) and our modified learning method for enhanced interpretability (**Fig. 4a**). Hidden node dropout strongly improved the correlation of node weights between network replicates (**Fig. 4a**, inset), resulting in lower variability of node weights and higher robustness of model interpretations (**Supplemental Fig. 7**). Indeed, learning with dropout resulted in much broader spreading of weights across the network and effectively prevented overreliance on few randomly selected inputs, as illustrated by an animation of the learning process (**Supplemental Video 1**). The gain in robustness was most pronounced between 0% and 10% dropout, and there was little impact on prediction performance up to a dropout rate of 30% (**Fig. 4a**, inset). Based on these observations, we selected a conservative dropout rate of 10% for further analysis, which resulted in a median node weight correlation of 0.912 (interquartile range 0.814 to 0.961), compared to 0.737 without dropout (interquartile range 0.608 to 0.854). Prediction performance was maintained with a median ROC AUC value of 0.982 with dropout (interquartile range 0.977 to 0.987), compared to 0.984 without dropout (interquartile range 0.979 to 0.987).

**Figure 4.**
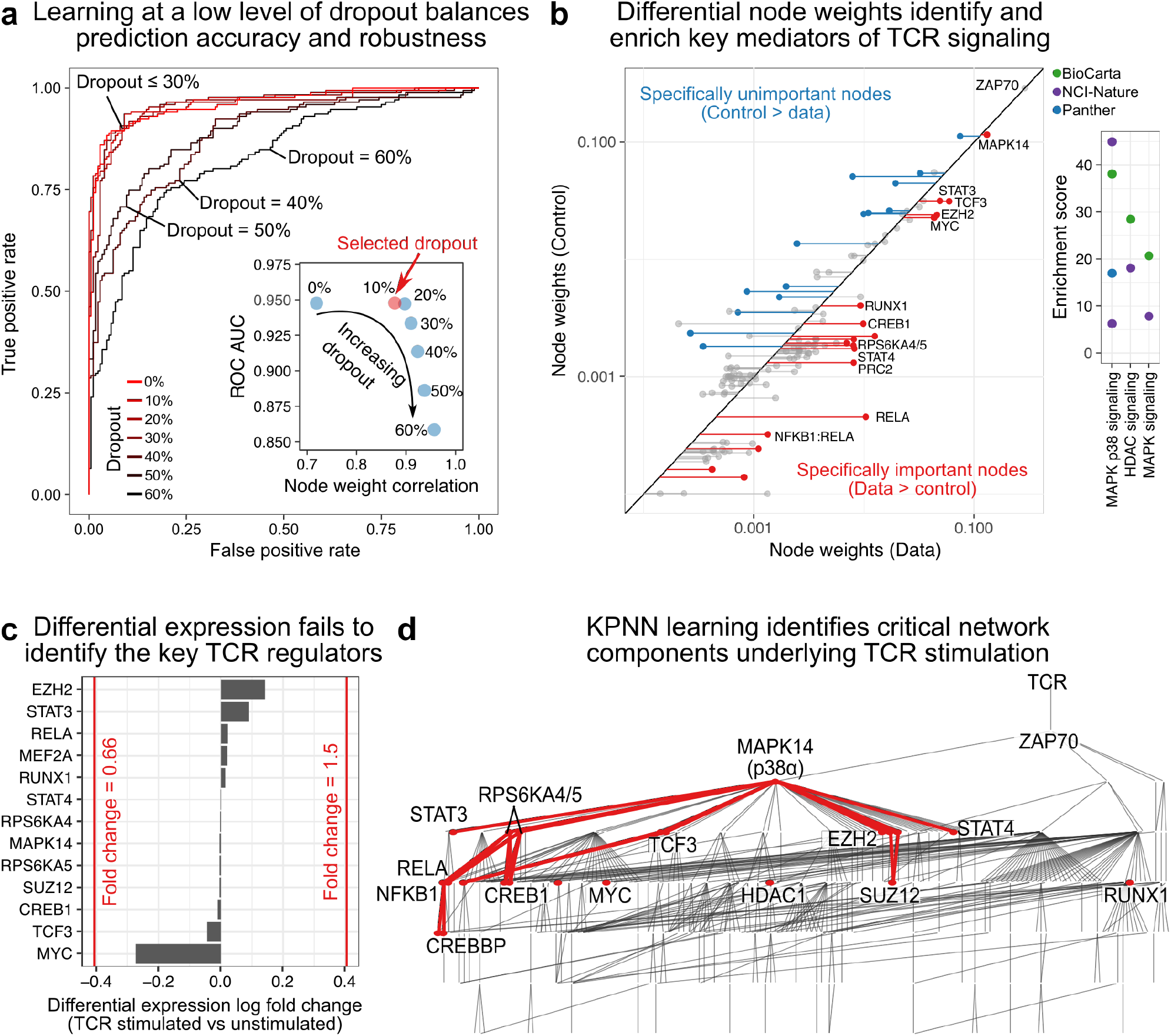
KPNN analysis of TCR stimulation. **(a)** Receiver operating characteristic (ROC) curves for the TCR KPNN predicting TCR stimulation with different levels of dropout. Curves for networks with ROC area under the curve (AUC) values closest to the mean obtained for each dropout rate are shown for illustration. The inset shows the mean ROC AUC values for different dropout rates (measuring prediction performance) and the mean correlation of network replicates (measuring network robustness). **(b)** Differential node weights at a dropout rate of 10%, comparing networks trained on actual data (x-axis) and on control inputs (y-axis). Nodes with p_adj_ below 0.05 are shown. (Right) Pathway enrichment among differential nodes. **(c)** Log fold change (LogFC) of gene expression for TCR regulators identified by the KPNN. Commonly used thresholds for differential expression (fold change = 1.5 and fold change = 0.66) are indicated in red. **(d)** Full TCR KPNN with the subnetwork of significantly differential nodes (p_adj_ < 0.05, at a dropout rate of 10%) highlighted in red.

Node weights were normalized for uneven connectivity in the TCR network by calculating the difference between the KPNN trained on the actual TCR stimulation dataset and the same KPNN trained on control inputs (**Fig. 4b, Supplemental Table 1**). The resulting differential node weights are directly interpretable as measures of protein/gene importance, and they indeed highlighted key regulators of TCR signaling, including the NF-κB1:RELA (p65:p50) transcription factor complex, its subunits RELA and NF-κB1, and the p38α MAP kinase MAPK14. Regulators with high differential node weights were generally enriched for MAP kinase pathways (**Supplemental Fig. 8, Supplemental Table 2**), which are known to be involved in TCR signaling and immune response^46^. Importantly, these regulators of TCR stimulation were not detected by analyzing their gene expression levels, which did not differ significantly between simulated and unstimulated cells (**Fig. 4c**). These results illustrate how deep learning on KPNNs quantifies regulatory activity that is not reflected in a regulator’s expression level but on the entire regulatory hierarchy upstream and downstream of the regulator.

The regulators of TCR signaling identified by our analysis formed a subnetwork within the KPNN (**Fig. 4d**). This subnetwork includes many regulators linked to TCR signaling, such as the transcription factors CREBBP, CREB1, and ATF^47,48^ as well as the kinases RPS6KA4 and RPS6KA5, which regulate transcription factors downstream of MAPK signaling. It further comprises the well-established T cell regulators STAT3^49^, STAT4^50^, HDAC1^51^, TCF3^52^, and RUNX1^53,54^. Finally, the subnetwork highlighted by the trained KPNN includes MYC^55^ and the PRC2 complex with subunits EZH2 and SUZ12^56,57^, which are general regulators of hematopoiesis and T cell development and expected to contribute to TCR-induced cell proliferation. Of note, ZAP70 and TCR were not part of this subnetwork – given their location at the apex of the KPNN, these two nodes received the highest possible node weights for both the actual data and the control inputs (**Fig. 4b**).

Importantly, the KPNN analysis of the TCR dataset reinforced and validated key design decisions of our modified learning method. First, dropout on hidden and input nodes dramatically enhanced the consistency of the learned node weights (**Fig. 4a**), and our chosen dropout rate of 10% resulted in the strongest enrichment for genes and proteins with a known role in TCR signaling (**Supplemental Fig. 8**). Second, normalization for the effect of uneven network connectivity was essential to obtain interpretable node weights, given that closeness to the TCR output node would otherwise dominate over the relevant biological information contained in the single-cell sequencing data (**Supplemental Fig. 9**).

In summary, interpretable deep learning using KPNNs uncovered a subnetwork of TCR-related regulators and signaling pathways that are relevant in this biological model of TCR stimulation. The identified subnetwork was extracted from a much broader network of potential regulators that were directly or indirectly linked to TCR signaling according to public databases. Importantly, the results depended on our modified learning method and could not have been obtained through classical gene expression analysis.

### KPNNs enable interpretable deep learning on massive-scale single-cell atlas data

To demonstrate the scalability of our method to a much larger dataset, we applied KPNNs to a Human Cell Atlas (HCA) reference dataset of 483,084 immune cells (T cells, B cells, and monocytes), derived from two sample sources (bone marrow and cord blood). While regulatory differences between these three cell types are well-established, here we focused on differences for the same cell types obtained from bone marrow versus cord blood. To that end, we trained multiclass KPNNs to predict cell type from gene expression, separately for each of these two sources of immune cells. From the trained KPNNs, we then extracted node weights reflecting the importance of multiple key regulator proteins in controlling gene expression of immune cells.

Our modified learning method achieved high prediction performance with a median ROC AUC value of 0.993 at a dropout rate of 20% (interquartile range 0.992 to 0.995), compared to 0.994 without dropout (interquartile range 0.992 to 0.995). Learning with dropout improved the robustness of the learned KPNNs (median node weight correlations: 0.703; interquartile range 0.652 to 0.751) compared to no dropout (0.593; interquartile range 0.509 to 0.670) (**Supplemental Fig. 10**). The trained KPNNs thus provide a solid foundation for analyzing and interpreting the regulatory mechanisms underlying immune cell identity in this cell atlas dataset.

Differential node weights (obtained by comparing trained KPNNs based on actual data versus control inputs) identified a core immune regulatory network that was shared across immune cell types, with certain cell-type-specific or tissue-specific elements (**Fig. 5a, Supplemental Table 3**). This network comprised transcription factors with a well-established role in immune regulation, including STAT5A^58,59^, STAT6^58,59^, JUN, CEBPB^60^, CREB1^61^, EP300^61^, and SP1^62^, as well as the tyrosine kinases ABL1 and GSK3B^63^, the cytokine receptor IL1R1, and the immune-regulatory RNA binding protein ILF3^64^. Furthermore, we identified important regulators of cell fate such as GATA1/2^65^, TAL1^66^, RUNX1^53,54^, EGR1^67^, ATF1^61^, EZH2^56,57^, MYC^55^, POU5F1^68^, SOX2^69^, and KLF4^70^, which are critically involved in the establishment of cell identity and hematopoietic development.

**Figure 5.**
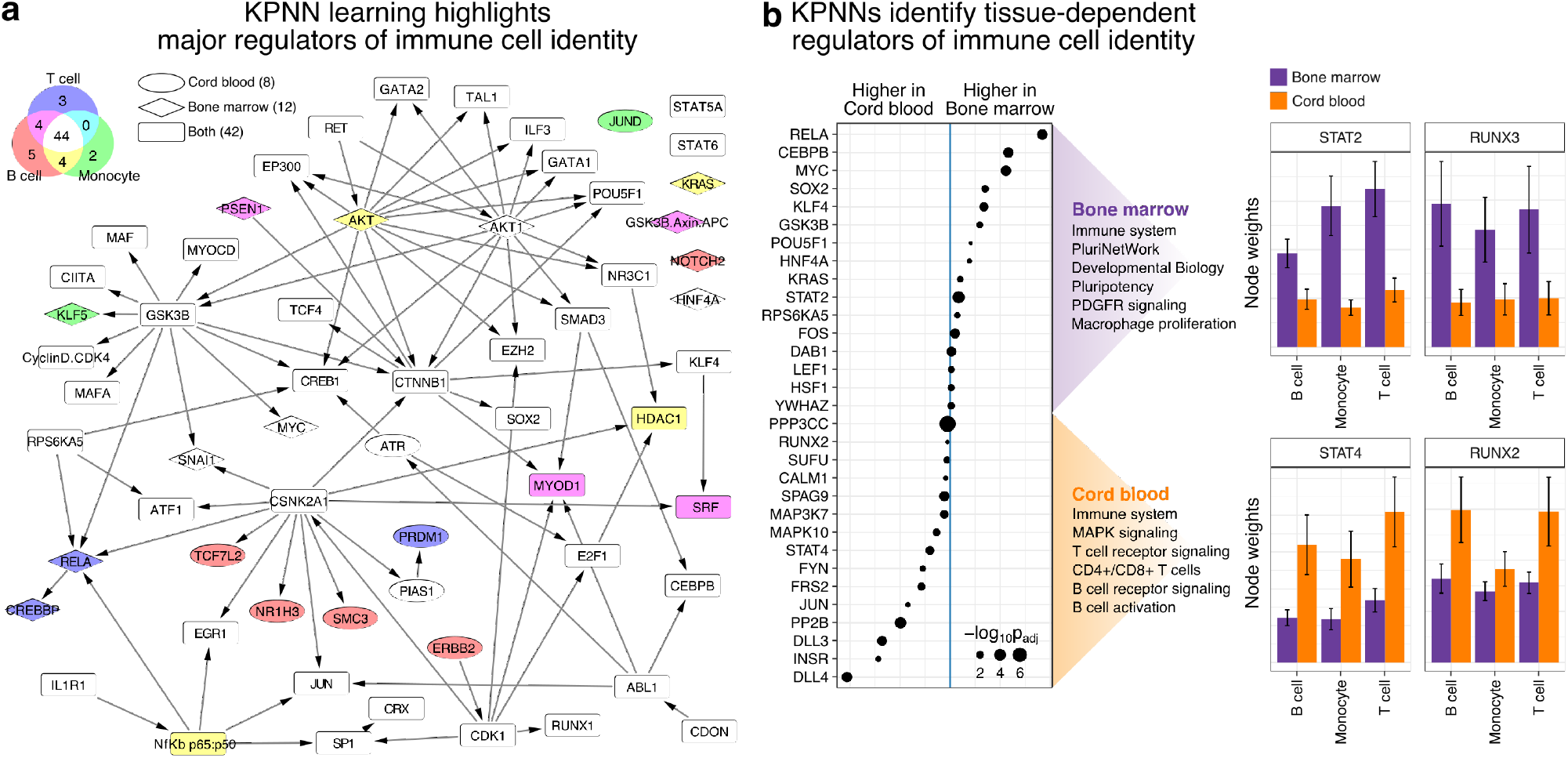
KPNN analysis of HCA immune reference maps. **(a)** Subnetwork of significantly differential nodes (p_adj_ < 0.05, at a dropout rate of 20%) from the HCA KPNN. Nodes are labeled based on the cell type (fill) and sample source (shape) where the nodes were identified as carrying differential weights between actual data and control inputs. **(b)** Comparative analysis of bone marrow and cord blood. (left) The plot shows the top-15 nodes with the most differential node weights when comparing bone marrow and cord blood. Gene set enrichments displayed on the right were calculated from the full list of nodes with differential weights. (right) Node weights of selected genes/proteins are shown for illustration. Error bars indicate standard error of the mean.

Based on the trained KPNNs, we sought to identify differences in immune cell composition and gene-regulatory mechanisms between bone marrow and cord blood. To this end, we compared node weights derived from KPNNs trained on single-cell transcriptome profiles of 262,895 immune cells from bone marrow and 220,189 immune cells from cord blood (see Methods for details). Nodes with higher weight for bone marrow comprised major regulators of cell fate such as SOX2, KLF5, KLF4, MYC, KRAS, and POU5F1, and we observed strong enrichment for related biological processes such as development, proliferation, and pluripotency (**Fig. 5b, Supplemental Fig. 11, Supplemental Tables 4 and 5**). In contrast, nodes with higher weight for cord blood comprised regulators associated with functions of mature immune cells, including multiple MAP kinases and protein phosphatases, and we observed enrichment for pathways such as T cell receptor signaling, B cell receptor signaling, B cell activation, and innate immune response.

Beyond this general tendency toward more developmentally immature regulatory profiles in bone marrow derived (as opposed to cord blood derived) B cells, T cells, and monocytes, we observed striking differences for individual regulators (**Fig. 5b**). First, the transcription factor STAT2 carried a much higher node weight in bone marrow, whereas the closely related STAT4 was more important in cord blood. Both proteins are part of the JAK-STAT signaling pathway, which mediates gene expression changes upon cytokine stimulation of immune cells^58,59^. STAT2 is well-known for its role in type I interferon signaling^59^, while STAT4 is primarily linked to IL-12 signaling and T helper cells^71^. Our results suggest that STAT2’s role in bone marrow derived cells is substituted in part by STAT4 in the more mature cord blood derived cells. Second, we observed a similar shift for RUNX3 (more important in bone marrow derived immune cells) versus RUNX2 (more important in cord blood derived immune cells). RUNX2 is primarily known for its role in osteoblasts and skeletal development^72^, but has also been implicated in thymic development and viral response of T cells^73,74^. RUNX3 is critical for T cell development^75,76^ and potentially more broadly relevant for hematopoietic development^77^. Our results corroborate the importance and developmental stage dependence of RUNX2 and RUNX3 in immune cell regulation.

In summary, we successfully performed interpretable deep learning using KPNNs on massive-scale singlecell atlas data, thereby validating the scalability of our method and establishing a broadly applicable approach for analyzing the single-cell compendia that are currently being produced for a broad range of human tissues. Comparing the regulatory processes underlying immune cells in the bone marrow and cord blood, we found strong evidence for developmentally immature B cells, T cells, and monocytes in bone marrow, as compared to cord blood. This observation is not due to differences among hematopoietic stem cells^78^, which were not included in our analysis. Rather, it captures relevant gene-regulatory differences among the bulk of B cells, T cells, and monocytes between these two immune cell compartments, likely reflecting cell-intrinsic differences as well as differences in cell-cell signaling and the tissue microenvironment.

## DISCUSSION

Deep learning has great potential for predictive analysis of biological phenomena, but is severely handicapped by its “black box” character. Even in those cases where deep neural networks already reach super-human prediction performance, they fail to provide human-understandable, high-level descriptions of the patterns they exploit for their successful predictions. To address this shortcoming, interpretable deep learning has recently emerged as an active area of machine learning research^44,79^. This research has focused almost exclusively on ex-post interpretation and reverse-engineering of learned models, for example by visualizing what ANNs trained for image recognition “see” when they classify a picture as a cat or a house^80–82^, or by inspecting DNA sequence motifs that are predictive of tissue-specific enhancer regions^31,83^. In the current study, we propose, implement, and validate a very different approach for interpretable deep learning in biology. We show that deep learning can be performed directly on biological networks (including signaling pathways and gene-regulatory networks), where each node corresponds to a gene or protein, and each edge corresponds to a regulatory interaction. This work thus demonstrates deep learning with ex-ante, built-in, molecular interpretability.

Several technologies converged to enable interpretable deep learning on biological networks using single-cell sequencing data. First, two decades of research into systems biology and regulatory networks have established large databases of signaling pathways and regulatory interactions, built on high-throughput experiments and on the curation of many individual mechanistic studies. Second, recent progress in single-cell sequencing makes it possible to obtain transcriptome profiles for thousands (millions) of single cells, thus providing sufficient experimental data for fitting complex multi-layer models and, more generally, the required scale for deep learning to play out its strengths. Third, deep learning has emerged as a method for inferring hidden states in deep neural networks based on large-scale training datasets, potentially allowing us to infer unobserved states in complex biological networks. By combining the breadth of potential regulatory interactions from public databases with large single-cell transcriptome data for the biological system under investigation, deep neural networks may indeed be able to pinpoint and connect those regulatory mechanisms that are relevant to the biological question of interest, thereby unraveling core regulatory networks and inferring regulatory cell states.

Our proof-of-concept for interpretable deep learning comprises two key components: Knowledge-primed neural networks (KPNNs) and an optimized learning method for enhanced interpretability. We derived application-specific KPNNs that model the regulatory processes of a cell by connecting cell surface receptors via signaling pathways and transcription factors (hidden nodes) to their target genes (input nodes corresponding to gene expression profiles obtained by single-cell RNA-seq). We demonstrated the feasibility of deep learning on these KPNNs and found that the prediction performance was comparable to that of ANNs. A comparison of network structures showed that KPNNs are very different from ANNs, with greater sparseness and higher modularity. The structural properties of KPNNs enable and enhance interpretability of trained models. First, all network elements in KPNNs have direct biological interpretations, with nodes representing genes/proteins and edges representing regulatory interactions. Second, sparsity restricts learning in KPNNs in that specific hidden nodes are connected to specific hidden nodes and inputs nodes based on biological knowledge. As a result, in a trained KPNN, highly weighted hidden nodes (proteins/genes) are key mediators for connecting the inputs (gene expression) with the output (cell type / cell state) and likely constitute relevant regulatory biology. This type of interpretability is absent from ANNs, where nodes are unspecifically connected with each other. It is also inaccessible to classical gene expression analysis because changes in the regulatory activity of key regulatory proteins often occur without concomitant changes in the expression of the corresponding gene.

To realize the full potential of KPNN interpretability, we modified the learning method in three ways, thereby addressing important conceptual and practical obstacles to interpretable deep learning. First, we temporarily removed a random subset of hidden nodes in each training step using a method called dropout^40^. This step conferred robustness to learned KPNN interpretations, in particular in the case of redundancy in signaling pathways. Second, we temporally removed random subsets of the input data during learning via dropout of input nodes, which enhanced the quantitative interpretability of node weights. Third, we trained the KPNNs on artificial control inputs in addition to the actual single-cell RNA-seq data, and we used the resulting control node weights to normalize the actual node weights for the uneven connectivity inherent to biological networks.

As a proof-of-concept for interpretable deep learning on biological networks, we applied our method to two single-cell sequencing datasets. Single-cell RNA-seq compendia are being produced at a rapidly growing rate, both within and beyond the Human Cell Atlas, which has led to an urgent need for bioinformatic methods that can derive biological insight and concrete molecular/mechanistic hypotheses from these valuable, large-scale datasets. Existing interpretive methods have focused primarily on differential gene expression and/or co-expression networks^84,85^. However, we observed that gene expression alone did not capture relevant changes in the regulatory activity of proteins. Proteins regulate gene expression in complex networks but their activity states remain hidden to measurements of gene expression. The activity of regulatory proteins thus constitutes hidden information similar to the hidden neurons of neural networks. Built on the powerful learning algorithms available for deep neural networks, KPNNs can learn hidden states of numerous regulatory proteins – without requiring genome-scale measurements of protein activity in single cells, which would be need for more classical mathematical modeling of signaling pathways and gene-regulatory networks^37,86,87^. As we demonstrated on simulated data with known ground truth, the node weights in the KPNNs indeed reflected protein regulatory activity in a quantitative manner, and they can be used in downstream analysis in much the same way as direct measurements of regulator protein activity, for example in differential analysis or gene set enrichment analysis.

In the first application, based on RNA-seq profiles of TCR stimulation *in vitro*, interpretable deep learning using KPNNs recapitulated much of what is known about TCR signaling, adapted to the concrete biological model (Jurkat cells stimulated by CD3/CD28 antibodies). From a broad regulatory network and application-specific gene expression data, KPNNs identified a dense subnetwork of TCR-related regulators linked to TCR signaling, including the transcription factors RELA and NF-kB1 as well as regulation by MAP kinases, and the more recently identified role of HDACs^51^. In the second application, a large HCA pilot dataset comprising 483,084 B cells, T cells, and monocytes obtained from bone marrow and cord blood, KPNNs identified a core network regulating immune cells. Comparing trained KPNNs between bone marrow and cord blood, we found that regulatory processes of immune cells in bone marrow were strongly associated with cellular development and proliferation, while processes in cord blood were associated with immune functions in more differentiated cells.

These two applications – together with our extensive analysis of simulated data – support the validity and practical utility of interpretable deep learning using KPNNs. Nevertheless, there are three potential limitations affecting our method that should be considered by its potential users. First, the sensitivity of the KPNNs (i.e., ability to detect the most relevant regulatory mechanisms for the biological system of interest) depends on the coverage of these mechanisms in the databases from which we derived our underlying biological, regulatory network. Fortunately, this is less of a limitation than it may seem, given that we include prior knowledge from any tissue or biological system, and rely on the learning process to confer specificity. Second, KPNNs constitute a simplification of biological networks, as they need to constitute directed acyclic graphs in order to be compatible with established deep learning algorithms. We have developed a procedure for obtaining high-confidence KPNNs from biological networks that include cycles, but there will likely be application that would benefit from more explicit modeling than KPNNs can currently provide (e.g., modeling of feedback loops). Third, the specificity of the KPNNs (i.e., ability to confidently exclude regulatory mechanisms that are not relevant in the biological system of interest) depends on the quality of the single-cell sequencing data, and noisy or biased experimental data may affect the performance of our method. We found that our method coped well with the inherently low coverage and frequent experimental dropouts in single-cell RNA-seq data, which was compensated by the large number of single-cell transcriptomes used as training data.

From a machine learning perspective, KPNNs complement graph neural networks (GNNs) in interesting ways. GNNs build on domain network knowledge to define relationships between input nodes, which are iteratively updated based on their neighbors prior to predicting a given output^88,89^. Typical applications of GNNs include predictions of node labels (given labels of some nodes) or graph labels (given a set of graphs). In biomedical research, GNNs have found initial application for the prediction of clinical attributes from gene expression^90^. While GNNs and KPNNs both build on prior network knowledge, GNNs use network knowledge to share information between inputs in order to improve prediction performance. Interpretability of GNNs is thus restricted to the (updated) inputs. In contrast, KPNNs use network knowledge to connect hidden nodes in order to achieve a deeper interpretability of the trained model. Toward this end, KPNNs exploit the analogy between biological regulation (signals are transduced from receptors via signaling proteins to transcription factors, which control gene expression) and feed-forward neural networks (output nodes, hidden nodes, input nodes), thus providing a conceptually novel way of performing deep learning using domain-specific graphs.

In conclusion, this study provides a general framework and initial proof-of-concept for interpretable deep learning on biological networks. We demonstrated the method’s utility for the molecular interpretation of single-cell RNA-seq data, which is a promising and exciting field of ongoing research. Moreover, given broad interest in networks for describing mechanistic processes that regulate a wide spectrum of biological entities, we expect that our use of deep learning on biological networks will be relevant also in other areas, for example analyzing metabolome/proteome data, biochemical reaction networks, cellular differentiation, or even brain circuits.

## METHODS

### Construction of a generic regulatory network for human cells

The knowledge-primed neural networks (KPNNs) used in this study connect the expression levels of individual genes via transcription factors and signaling proteins to the cell surface receptors to model cell signaling pathways and their cross-talk in one network. As the biological basis of the KPNNs, we constructed a generic regulatory network by integrating transcription factor and signaling pathway data from multiple public databases into a comprehensive model of the potential regulatory space in human cells.

#### Transcription factors

To connect the genes whose expression is measured by single-cell RNA-seq (they constitute the input nodes of the KPNNs) to the transcription factors that may regulate their activity (they constitute part of the KPNN’s hidden nodes), we obtained transcription factor / target gene pairs from the Harmonizome^91^ database (as of 3 July 2017), including the following data sources: ENCODE, CHEA, ESCAPE, MotifMap, and TRANSFAC (manually curated and predicted). Further transcription factor / target gene pairs were obtained from the TTRUST^92^ database (as of 3 July 2017), which we mapped to gene names using mapping data from NCBI (ftp://ftp.ncbi.nlm.nih.gov/gene/DATA/gene_info.gz, as of 10 October 2016). These datasets were then combined by counting the number of datasets that support a connection for each transcription factor / target gene pair. To prioritize experimental over computational evidence, weights of 1.0 and 0.3 were used for experimental and computational support, respectively. Then, for each gene, the transcription factors with the largest weighted number of datasets supporting the connection to the gene were retained. As a result of this procedure, 745 genes were targeted by more than 25 transcription factors; these genes were removed from the analysis to avoid biasing the network with promiscuous or false-positive regulatory interactions.

#### Signaling pathways

To connect signaling proteins (they constitute part of the KPNN’s hidden nodes) to their target proteins (transcription factors or other signaling proteins), we obtained a comprehensive dataset of protein signaling interactions, protein complexes, and protein family information from the SIGNOR^93^ database (as of 3 July 2017), a large and manually curated database of directed signaling interactions. To focus this network on signaling pathways that connect proteins, we removed interactions of transcriptional regulation, guanine nucleotide exchange factor, transcriptional activation, post-transcriptional regulation, or transcriptional repression; and we removed all nodes that were not proteins, complexes, or protein families, as well as nodes from databases other than UniProt^94^ and SIGNOR. All retained interactions were then aggregated into a directed graph of potential regulatory interactions. To capture interactions between complex members and annotations of protein families, we further linked family members to nodes representing protein families, and protein complex members to nodes representing complexes. This was done using bidirectional edges, thus enabling protein complex members to regulate their target complexes and vice versa.

### Construction of application-specific KPNNs

From the generic regulatory network described in the previous section, we derived application-specific KPNNs as follows: (i) we defined the cell surface receptors expected to be most relevant for the biological phenomenon of interest (default: all cell surface receptors included in the network); (ii) we extracted a directed acyclic graph that connects the receptors to all reachable transcription factors; and (iii) we connected transcription factors to their target genes. The resulting graphs were then used for interpretable deep learning by reversing the cascade: Measured gene expression values (input nodes) predicted the activity states of connected transcription factors (hidden nodes), which predicted the activity states of signaling proteins (hidden nodes), which then predicted the activity states of surface receptors. In the TCR KPNN, the TCR is the output node that predicts stimulation. In the HCA KPNN, receptors are hidden nodes that ultimately predict cell type (output nodes).

#### TCR KPNN

To construct the KPNN for TCR signaling, we selected the TCR node (SIGNOR-C153) as the output node of the network, and we calculated shortest paths to all reachable transcription factors in the generic regulatory network using the function *all_shortest_paths* from the igraph package (version 1.1.2) in R (version 3.2.3). These paths were then combined, resulting in a directed acyclic graph. Finally, transcription factor / target gene pairs were used to connect each transcription factor to its target genes (input nodes).

#### HCA KPNN

To generate the KPNN predicting cell type in the Human Cell Atlas (HCA) reference dataset, three nodes that denote the three cell types were added to the generic regulatory network and connected to all cell surface receptors, thus modeling cellular regulation of gene expression as the ensemble of signaling pathways. Cell surface receptors were identified based on the pathway annotations in the SIGNOR database using SIGNOR’s REST API. We then calculated shortest paths from the output nodes via the cell surface receptor nodes to all reachable transcription factors in the generic regulatory network using the function *all_shortest_paths* from the igraph package. These paths were combined, resulting in a directed acyclic graph. Transcription factor / target gene pairs were used to connect each the transcription factors to their target genes (input nodes). This initial graph was then extended to ensure inclusion of all relevant cell surface receptors. To do so, shortest paths were iteratively added from individual receptors to all reachable transcription factors under the condition that they do not introduce cycles. This was ensured by to the following procedure: (1) Shortest paths from all receptors to all transcription factors were identified in the generic regulatory network and transformed to an edge list. (2) Edges from step 1 were added to the graph if the parent depth was smaller than the child depth, or if the child was not yet included in the network. Node depth was defined as the distance of a node from the output nodes. In addition, to rule out feedback loops from transcription factors to upstream protein signaling pathways, the depth of transcription factors (and their downstream nodes) was artificially set to be greater than all non-transcription factors. Finally, step 1 and 2 were repeated iteratively until no more edges could be added. This procedure thus enabled us to extend our graph by adding relevant connections, while ensuring that we did not introduce cycles to the graph.

### Single-cell transcriptome datasets used for KPNN training

While the KPNNs provide comprehensive, knowledge-based models of the wider regulatory space that could potentially be relevant and interpretable in the application of interest, it is the deep learning on KPNNs using single-cell transcriptome data that confers specificity and identifies those parts of the KPNNs that are indeed relevant and predictive for the concrete biological application. In this study, we analyzed two real-world single-cell transcriptome datasets (TCR, HCA) as well as a set of simulated transcriptome profiles with known ground truth, which we used for methods development, validation, and benchmarking.

#### TCR dataset

The TCR dataset was downloaded from GEO (GSE137554). The dataset is based on the CROP-seq assay^41^ combined with single-cell RNA-seq using the 10x Genomics platform. Only unperturbed cells were used in the analyses for this paper. To identify the CRISPR guide RNAs in each cell, a reference genome was created by extending the human GRCh38 genome assembly with sequences of all used guide RNAs with Cell Ranger *mkref*. Sequencing data were aligned to this reference genome with Cell Ranger *count* and merged with Cell Ranger *aggr*. Barcodes with less than 500 unique molecular identifiers (UMIs), less than 200 identified genes, or more than 15% mitochondrial reads were considered low-quality and removed from the analysis. Cells expressing only non-targeting guide RNAs were selected for further analysis. Class values for TCR stimulation were assigned based on whether a cell was part of the TCR stimulated or unstimulated sample.

#### HCA dataset

The HCA dataset was downloaded from the “Census of Immune Cells” that is part of the Human Cell Atlas (https://preview.data.humancellatlas.org/), as of 31 July 2018. Low-quality cells with less than 500 unique molecular identifiers (UMIs), less than 200 identified genes, or more than 15% mitochondrial reads were removed from the analysis, and expression levels were transformed to log(TPM + 1) values. Cell types (class values) were assigned based on the expression of cell-type-specific marker genes. Marker gene expression for CD79A and CD19 was used to identify B cells; CD3D, CD3G, and IL32 to identify T cells; CD14 and CST3 to identify monocytes. Cells with log(TPM + 1) values above 4 for any of the listed marker gene were assigned to the respective cell types. Cells assigned to none of the cell types (which comprises all other cell types) or to multiple cell types (including cell duplicates) were removed from the analysis. Using this procedure, 483,084 from a total of 628,630 cells were uniquely assigned to one of the three cell types.

#### Simulated data

Data with a defined ground truth were simulated based on the TCR dataset. We averaged the expression counts for the unstimulated state for each gene to generate a baseline gene expression profile. Next, a positive or negative twofold change was introduced in selected genes (predictive/informative genes, whose expression was linearly correlated with each cell’s class value), resulting in a second (differential) gene expression profile. Finally, single-cell RNA-seq profiles of individual cells were simulated by subsampling reads from these average profiles using *rbinom* in R as described previously^95^. 2,000 single-cell RNA-seq profiles were generated, 50% of which were derived from the baseline profile and 50% from the differential profile.

### Deep learning methodology and implementation

Interpretable deep learning was performed on the application-specific KPNN with the corresponding single-cell transcriptome profiles as training data, using a three-step workflow: (i) processing of input data, (ii) training of the KPNN as a deep neural network, and (iii) evaluation of prediction performance on unseen test data. This workflow was implemented in a custom Python (version 2.7.6) program, using the Python libraries tensorflow^96^ (version 1.3.1), pandas (version 0.19.2), scipy (version 0.14.0), and numpy (1.13.2). This program accepts three inputs: (i) a neural network graph (KPNN or ANN) in the form of an edge list, (ii) a file containing class values for each single cell, and (iii) a file containing transcriptome profiles for each single cell.

#### Processing of input data

Input data were split into training set (60% of samples), validation set (20%), and test set (20%) using numpy *random.choice*. Gene expression data were converted to log(TPM + 1) values and normalized for each gene to a maximum value of one and a minimum value of zero. Normalization factors were calculated based on the training data and then applied to the validation and test data. For the HCA dataset, minibatches were obtained using numpy *random.shuffle*.

#### Network training

Training of the neural network requires initialization of edge weights, an activation function for nodes, a loss function, an algorithm to minimize the loss function, and a process to terminate training. These elements were implemented using tensorflow, with the following setup. Edge weights were randomly initialized using tensorflow *global_variables_initializer*. A sigmoid activation function was used for all neurons. The loss function to minimize was chosen as a weighed cross-entropy with L2 regularization. To calculate the loss function, cross-entropy was first calculated using tensorflow *nn.sigmoid_cross_entropy_with_logits*. Then, to improve training in the presence of class imbalance, the cross-entropy of each sample was weighted by the number of samples of each class. The weight for each class was calculated as (1 / N_classes_) / (N_x_ / N_samples_), where N_classes_ is the number of classes, N_x_ is the number of samples of class x, and N_samples_ is the total number of samples. Finally, L2 regularization was calculated using tensorflow *nn.l2_loss* and added to the weighted cross-entropy. Using this loss function, the ADAM algorithm was used to minimize the regularized, weighted cross-entropy using tensorflow *train.AdamOptimizer*. To track learning progress, training set and validation set error was calculated by subtracting predicted class probabilities from true labels. Training was terminated using early stopping. Specifically, epochs were considered failed when either training loss plateaued (indicative of arrival at a minimum) or validation set error increased (indicative of overfitting), and learning was stopped once a specified number of failed epochs was reached. To be able to recover the most promising model, new models were saved during training (using tensorflow *train.Saver*) if they decreased validation set error by a given percentage compared to the latest saved model. When a model was saved, the counter of failed epochs was reset to zero. After learning was terminated, the most recently saved model was loaded.

#### Test data performance

Using the latest saved model, the error on the (previously unseen) test set was assessed and receiver-operator characteristic curves were derived as the final performance metric.

#### Hyperparameters

Training hyperparameters were chosen for each dataset by inspecting the learning progress using tensorflow’s TensorBoard, which enables live tracking of training loss, validation error, validation loss, and edge weights. The two parameters used for early stopping (minimum percent improvement on the validation set error required to save a model; number of allowed failed learning epochs) were chosen to minimize the number of iterations run in the plateau of the learning curve. Learning rate (alpha) was selected to the highest value that ensured smoothness of learning curves. L2-loss regularization parameter (lambda) was chosen at the largest value that did not result in weights shrinking to zero.

### Optimized learning method to enable interpretability

On top of the standard deep learning methodology (as described in the previous section), three modifications were implemented in order to enable and enhance interpretability of the trained KPNNs: (i) Dropout on hidden nodes to improve robustness, (ii) dropout on input nodes to increase quantitative interpretability, and (iii) training on simulated control inputs to normalize for uneven connectivity in biological networks.

#### Dropout on hidden nodes and on input nodes

Dropout constitutes a modification of the learning algorithm where a given percentage of nodes is randomly set to zero in each training iteration and for each sample, temporarily removing the contribution of the affected nodes. Dropout was originally developed as a regularization technique to reduce overfitting^40^. Applied to KPNNs, the removal of random nodes by dropout encourages networks to spread weights across all relevant nodes, which reduces variability of node weights across network replicates (leading to improved robustness) and balances weights across input nodes (leading to node weights that more quantitatively reflect node importance). Dropout was implemented in our network training program using tensorflow *nn.dropout*, which was applied separately to hidden nodes as well as input nodes.

#### Normalizing for the uneven connectivity of biological networks

To normalize for the inherent network structure and connectivity patterns of each KPNN, we trained the KPNNs not only on the actual single-cell transcriptome data, but also on simulated control inputs where all input nodes were artificially set to values that are equally predictive of the class values (linearly correlated with the output). All input nodes carry equal importance in this scenario, hence the resulting node weights reflect only the inherent network structure of the KPNN. This control analysis enabled quantification of, and normalization for, uneven connectivity in each KPNN. Control inputs were generated in the same way as the simulated single-cell transcriptome datasets used to develop our methodology (as described above), with one difference: While only a subset of inputs were selected as predictive in the simulated dataset, the control inputs were simulated such that all inputs were equally predictive. To that end, raw read counts were summed up across all cells corresponding to one class value (e.g., unstimulated cells in the TCR data) to generate an average transcriptome profile. A positive or negative (chosen with numpy *random.choice*) twofold change was added to all input genes to generate a second average transcriptome profile. Reads were then drawn from the two average transcriptome profiles using numpy *ran-dom.binomial*^95^. The number of reads drawn was based on the number of reads in each cell in the original data. For the TCR data and for the simulated data, the same number of cells as in the original dataset were simulated with control inputs. In the HCA analysis, 2,000 cells were simulated with control inputs.

### Training KPNNs for the TCR, HCA, and simulated datasets

Using the optimized KPNN learning method described in the previous section, we trained neural networks to predict class values from input data for our two real-world datasets (TCR, HCA) as well as the simulated transcriptome profiles with known ground truth. Trained KPNNs were subsequently analyzed to obtain biological interpretations. This section describes the parameters chosen to train KPNNs for each dataset.

#### TCR dataset

Raw reads of all 1,735 cells, class values (stimulated and unstimulated), and an edge list encoding the TCR KPNN structure were provided as input to the learning program, where they were normalized and processed as described above. Hyperparameters (alpha: 0.01; lambda: 0.1) were chosen based on the inspection of learning curves using TensorBoard. Learning was stopped after 20 failed epochs. During learning, new models were saved if they reduced the validation set error of the latest stored (i.e., so far best-performing) model by at least 20%. Dropout rates from 0% to 50% were evaluated in steps of 10 percentage points. In addition, dropout was adjusted for each node based on the number of parent nodes to account for the very sparse structure of the TCR KPNN, where networks failed to converge otherwise. Specifically, nodes with only one parent were never dropped, dropout was limited to 10% for nodes with two parents, and to 30% for nodes with three parents. Node weights were normalized based on the same KPNN trained on control inputs with dropout levels corresponding to the levels used for the original data.

#### HCA dataset

Raw reads of all 483,084 cells, class values (B cells, T cells, and monocytes), and an edge list encoding the HCA KPNN were provided as input to the learning program. Marker genes used for cell type assignment were removed prior to training. Hyperparameters (alpha: 0.05; lambda: 0.1) were chosen based on the inspection of learning curves using TensorBoard. Learning was stopped after 10 failed epochs. During learning, models were saved if they reduced the validation set error of the latest stored (i.e., so far best-performing) model by at least 30%. Dropout rates from 0% to 40% were evaluated in steps of 20 percentage points. Learning was performed in minibatches of 1,000 single cells. Node weights were normalized based on the same KPNN trained on control inputs with dropout levels corresponding to the levels used for the original data. Differences in the parameters used to train KPNNs on the HCA data and on the TCR data were due to the large difference in sample size (greater than two orders of magnitude), which resulted in improved learning progress per epoch but a much longer duration of each epoch for the HCA data.

#### Simulated dataset

Simulated raw reads, class values (0 and 1), and the edge lists of the simulated networks were provided as input to the learning program. Hyperparameters were selected as follows: alpha of 0.001 and lambda of 0.2 in networks containing only one predictive node; alpha of 0.05 and lambda of 0.1 in networks with two predictive nodes; alpha of 0.05 and lambda of 0.2 in networks with three weakly and three strongly predictive nodes; and alpha of 0.05 and lambda of 0.2 for the comparison of networks with and without control inputs. The dropout rate was set to 30% in all analyses where dropout was used. Learning was stopped after 20 failed epochs, and models were saved if they reduced the validation set error of the latest stored (i.e., so far best-performing) model by at least 20%.

### Calculation of node weights as a measure of node importance

Once KPNNs were trained, they were analyzed by calculating node weights as a reflection of node importance for the KPNN-based predictions. Edge weights are readily obtainable from the network model, but they only indicate the (local) relationship of a given node to its parent node and do not capture the (global) importance of each node for the full network. To obtain informative node weights, we applied small perturbations to all hidden nodes and measured changes in network output, thus quantifying the importance of each node to the output of the entire network. Finally, because the sign of the resulting node weights is largely arbitrary, we take the absolute value of the node weights as a measure of the importance of each node in the trained KPNNs.

#### Calculation of node weights using induced perturbations

To obtain informative node weights, we applied a procedure analogous to numerical gradient estimation^97^, which is commonly used to evaluate the effect of small numerical perturbations of edge weights on the network predictions (outputs) in order to test the calculation of backpropagation gradients. Here, instead of perturbing edge weights, we applied perturbations to nodes, thus estimating the effect that small perturbations of node outputs (activations) have on the network predictions. For each node ni, network predictions (class probabilities) were calculated after perturbing ni twice, once by adding and once by subtracting a small factor (epsilon, which was set to 0.001 for all datasets) from the output (activation) of ni. The mean difference of class probabilities after additive and subtractive perturbation was then further divided by 2*epsilon to derive a normalized measure of node weight. As a result, nodes with large node weights had a greater effect on network predictions than nodes with smaller node weights, thus quantifying the importance of each hidden node in the network.

#### The rationale for calculating absolute values of the node weights

Node outputs can impact network predictions positively or negatively, depending on the sign of their associated edge weights. In the case of input nodes, the sign of node weights can be interpreted directly with respect to network predictions (for example “high expression of gene X predicts TCR stimulation and low expression predict no stimulation”). For hidden nodes, however, this relationship is absent because hidden nodes can assign negative or positive weights to the associated input nodes with equal outcome. For example, a negative edge weight associated with a negative node output will result in a positive number as much as a positive edge weight associated with a positive node output. The sign of node weights will thus randomly differ between replicate networks (i.e., the same network trained separately on the same input data). In contrast, the absolute magnitude of node weights reflects the importance of each node for prediction, independent of its sign. For this reason, we calculated absolute node weights, which robustly quantify the contribution of each node to the predictions.

### Statistical analysis of node weights

After training of each KPNN, node weights were calculated as described in the previous section and exported for downstream analysis. Node weights were thus obtained from each replicate network. To perform comparisons between actual input data and simulated control inputs (and between bone marrow and cord blood for the HCA data) we performed statistical differential analysis on node weights in much the same way as it is commonly done for gene expression data. To this end, node weights were imported into R (version 3.2.3), where they were analyzed and plotted using packages data.table (version 1.11.4), ggplot2 (version 2.2.1), and pheatmap (version 1.0.10). Pearson correlations of node weights were calculated using the function *cor* in R.

#### Analysis of node weights for the TCR KPNN

Differential node weights were calculated by comparing node weights from actual data versus control inputs, and evaluated using gene set enrichment analysis. For these analyses, only highly predictive networks with test set error lower than 0.2 were retained. Networks with dropout greater than 30% were excluded due to their weak prediction performance. To avoid biases arising from sample size differences, the number of replicate networks in each group was downsampled to that of the smallest group (n = 42). Node weights of all networks were normalized using limma *normalizeQuantiles* in R. Significance of differential node weights was tested for each hidden node using the *t.test* function in R. P-values were corrected for multiple testing using the function *p.adjust* with parameter “BH”. Nodes with adjusted p-values below 0.05 were selected as significant. Gene set enrichment for significant genes/proteins was performed using Enrichr^98^, with databases *Panther_2016, NCI-Nature_2016*, and *BioCarta_2016*. In addition, to quantify the enrichment of annotated TCR signaling proteins, we used Enrichr to download all genes annotated with “TCR Signaling Pathway_Homo sapiens_WP69” from *WikiPathways_2016* as well as genes annotated with “TCR signaling in naive CD8+ T cells_Homo sapiens” from *NCI-Nature_2016* (December 15, 2017).

#### Analysis of node weights for the HCA KPNN

Differential node weights were calculated by comparing node weights from actual data versus control inputs (for the first analysis), and by comparing node weights between networks trained on bone marrow versus cord blood (for the second analysis). In both cases, only highly predictive networks with precision greater than 0.9 for all three cell types were used to obtain node weights. Similar to the TCR analysis, the number of replicate networks in each group was subsampled to that of the smallest group (n = 48) to avoid biases arising from sample size differences. Differential node weights relative to control input networks were calculated using *t.test*, separately for each cell type in bone marrow and cord blood. P-values were corrected for multiple testing using the function *p.adjust* with parameter “BH”. Nodes with adjusted p-value below 0.05 were selected as significant. For the comparison of bone marrow versus cord blood, differential analysis was performed with a linear model using the function *lm* in R. Coefficients were trained for cell type and for bone marrow versus cord blood. Coefficients and p-values for the comparison between bone marrow and cord blood were extracted from the fitted linear model using the R functions *coef* and *summary*. P-values were adjusted for multiple testing as described above, with a significance threshold of 0.05. Enrichment analyses of significant nodes were performed using Enrichr^98^, with the databases *WikiPathways_2016, Reactome_2016, Jensen_ TISSUES, KEGG_2016, NCI-Nature_2016, Panther_2016, BioCarta_2016*.

#### Analysis of node weights for the simulated data

Highly predictive networks with test set error below 0.1 were used to obtain node weights. Node weights were scaled to a minimum of 0 and maximum of 1 prior to plotting.

### Construction of generic ANNs and structural network comparison

To empirically define the characteristics of KPNNs, we compared and benchmarked them against generic artificial neural networks (ANNs). These ANNs were constructed such that they match the corresponding KPNNs in terms of the number of input nodes, hidden nodes, and output node(s). In contrast, the distribution of edges was notably different, and the number of hidden layers was handled as a free parameter ranging from 1 to the maximum depth of the corresponding KPNN, thus resulting in multiple ANNs for each KPNN.

#### Construction of ANNs

Fully connected ANNs (fANNs) were constructed by distributing the hidden nodes equally across all layers and adding edges to fully connect adjacent layers. Moreover, all nodes of the first hidden layer were connected to all input nodes, and all nodes of the last hidden layer were connected to the output node(s). Given that these fANNs have many more edges than their corresponding KPNNs, sparse ANNs (sANNs) were derived from the fANNs by down-sampling of edges, in order to reach the same number of edges and nodes as in the corresponding KPNN. This was done in two steps: First, we generated a minimal network that spans all nodes. To do so, edges were removed such that every node was connected to exactly one randomly selected node in the following layer and at least one randomly selected node in the previous layer. Second, edges were added back randomly until the network had the same number of edges as the corresponding KPNN. This step was carried out separately for the input layer and for the hidden layer of the network. As a result, the resulting sANN was equivalent to the KPNN in the number of edges connecting input nodes to hidden nodes, and also in the number of edges connecting hidden nodes amongst each other.

#### Analysis of network structure

Structural network analysis comparing KPNNs and ANNs was performed using the igraph package in R. Outdegree was measured with the igraph function *degree*. Distances to input nodes was measured with the igraph function *distances*. The resulting distance values were used to calculate reachability, counting the number of input nodes with finite distance. To compare modularity, the network was transformed into an undirected graph using the igraph function *as.undirected*, edges to remove were identified using the igraph function *edge.betweenness.community*, and the number of clusters was determined with the igraph function *clusters*.

### Data and code availability

All datasets are publicly available. The source code to train and analyze KPNNs is available as a GitHub repository https://github.com/epigen/KPNN.

## ACKNOWLEDGMENTS

The authors thank Patricia Carey for HPC cluster maintenance; Jörg Menche, Sylvia Knapp, Wilfried Ellmeier, Murat Tugrul, Lukas Folkman, Eugenia Iofinova, Stephan Reichl, Thomas Krausgruber, and Matthias Farlik for insightful discussions; and various other members of the Bock lab and of CeMM for their help and advice. This work was funded in part by an Austrian Science Fund (FWF) Special Research Programme grant to C.B. (FWF SFB F 6102). N.F. is supported by a fellowship from the European Molecular Biology Organization (EMBO ALTF 241-2017). C.B. is supported by a New Frontiers Group award of the Austrian Academy of Sciences and by an ERC Starting Grant (European Union’s Horizon 2020 research and innovation programme, grant agreement n° 679146).

## AUTHOR CONTRIBUTIONS

NF and CB designed the study; NF performed computational analyses; CB supervised the research; NF and CB wrote the manuscript.

## REFERENCES

1. Krizhevsky, A., Sutskever, I. & Hinton, G. E. ImageNet Classification with Deep Convolutional Neural Networks. in Advances in Neural Information Processing Systems 25 (eds. Pereira, F., Burges, C. J. C., Bottou, L. & Weinberger, K. Q.) 1097–1105 (Curran Associates, Inc., 2012).

2. Farabet, C., Couprie, C., Najman, L. & LeCun, Y. Learning Hierarchical Features for Scene Labeling. IEEE Trans. Pattern Anal. Mach. Intell. 35, 1915–1929 (2013).

3. Szegedy, C. et al. Going Deeper With Convolutions. in Proceedings of the IEEE Conference on Computer Vision and Pattern Recognition 1–9 (2015).

4. Hinton, G. et al. Deep Neural Networks for Acoustic Modeling in Speech Recognition: The Shared Views of Four Research Groups. IEEE Signal Process. Mag. 29, 82–97 (2012).

5. Graves, A., Mohamed, A. & Hinton, G. Speech recognition with deep recurrent neural networks. in 2013 IEEE International Conference on Acoustics, Speech and Signal Processing 6645–6649 (IEEE, 2013). doi:10.1109/ICASSP.2013.6638947

6. Collobert, R. et al. Natural Language Processing (Almost) from Scratch. J. Mach. Learn. Res. 12, 2493–2537 (2011).

7. Jean, S., Cho, K., Memisevic, R. & Bengio, Y. On Using Very Large Target Vocabulary for Neural Machine Translation. ArXiv14122007 Cs (2014).

8. Sutskever, I., Vinyals, O. & Le, Q. V. Sequence to Sequence Learning with Neural Networks. in Advances in Neural Information Processing Systems 27 (eds. Ghahramani, Z., Welling, M., Cortes, C., Lawrence, N. D. & Weinberger, K. Q.) 3104–3112 (Curran Associates, Inc., 2014).

9. Cho, K. et al. Learning Phrase Representations using RNN Encoder-Decoder for Statistical Machine Translation. ArXiv14061078 Cs Stat (2014).

10. Bahdanau, D., Cho, K. & Bengio, Y. Neural Machine Translation by Jointly Learning to Align and Translate. ArXiv14090473 Cs Stat (2014).

11. Mnih, V. et al. Playing Atari with Deep Reinforcement Learning. ArXiv13125602 Cs (2013).

12. Silver, D. et al. Mastering the game of Go without human knowledge. Nature 550, 354–359 (2017).

13. Silver, D. et al. A general reinforcement learning algorithm that masters chess, shogi, and Go through self-play. Science 362, 1140–1144 (2018).

14. Chen, C., Seff, A., Kornhauser, A. & Xiao, J. DeepDriving: Learning Affordance for Direct Perception in Autonomous Driving. in Proceedings of the IEEE International Conference on Computer Vision 2722–2730 (2015).

15. Huval, B. et al. An Empirical Evaluation of Deep Learning on Highway Driving. ArXiv150401716 Cs (2015).

16. Angermueller, C., Pärnamaa, T., Parts, L. & Stegle, O. Deep learning for computational biology. Mol. Syst. Biol. 12, 878 (2016).

17. Wainberg, M., Merico, D., Delong, A. & Frey, B. J. Deep learning in biomedicine. Nat. Biotechnol. 36, 829–838 (2018).

18. Ching, T. et al. Opportunities and obstacles for deep learning in biology and medicine. J. R. Soc. Interface 15, 20170387 (2018).

19. Eraslan, G., Avsec, Ž., Gagneur, J. & Theis, F. J. Deep learning: new computational modelling techniques for genomics. Nat. Rev. Genet. 1 (2019). doi:10.1038/s41576-019-0122-6

20. Esteva, A. et al. Dermatologist-level classification of skin cancer with deep neural networks. Nature 542, 115–118 (2017).

21. Helmstaedter, M. et al. Connectomic reconstruction of the inner plexiform layer in the mouse retina. Nature 500, 168–174 (2013).

22. Angermueller, C., Lee, H. J., Reik, W. & Stegle, O. DeepCpG: accurate prediction of single-cell DNA methylation states using deep learning. Genome Biol. 18, 67 (2017).

23. Alipanahi, B., Delong, A., Weirauch, M. T. & Frey, B. J. Predicting the sequence specificities of DNA- and RNA-binding proteins by deep learning. Nat. Biotechnol. 33, 831–838 (2015).

24. Leung, M. K. K., Xiong, H. Y., Lee, L. J. & Frey, B. J. Deep learning of the tissue-regulated splicing code. Bioinformatics 30, i121–i129 (2014).

25. Xiong, H. Y. et al. The human splicing code reveals new insights into the genetic determinants of disease. Science 347, 1254806 (2015).

26. Eraslan, G., Simon, L. M., Mircea, M., Mueller, N. S. & Theis, F. J. Single cell RNA-seq denoising using a deep count autoencoder. bioRxiv 300681 (2018). doi:10.1101/300681

27. Ding, J., Condon, A. & Shah, S. P. Interpretable dimensionality reduction of single cell transcriptome data with deep generative models. Nat. Commun. 9, 2002 (2018).

28. Rashid, S., Shah, S., Bar-Joseph, Z. & Pandya, R. Project Dhaka: Variational Autoencoder for Unmasking Tumor Heterogeneity from Single Cell Genomic Data. bioRxiv 183863 (2018). doi:10.1101/183863

29. Deng, Y., Bao, F., Dai, Q., Wu, L. & Altschuler, S. Massive single-cell RNA-seq analysis and imputation via deep learning. bioRxiv 315556 (2018). doi:10.1101/315556

30. Lopez, R., Regier, J., Cole, M. B., Jordan, M. I. & Yosef, N. Deep generative modeling for single-cell transcriptomics. Nat. Methods 15, 1053 (2018).

31. Kelley, D. R., Snoek, J. & Rinn, J. L. Basset: learning the regulatory code of the accessible genome with deep convolutional neural networks. Genome Res. 26, 990–999 (2016).

32. Cusanovich, D. A. et al. A Single-Cell Atlas of In Vivo Mammalian Chromatin Accessibility. Cell 174, 1309–1324.e18 (2018).

33. Tan, J. et al. Unsupervised Extraction of Stable Expression Signatures from Public Compendia with an Ensemble of Neural Networks. Cell Syst. 5, 63–71.e6 (2017).

34. Lin, C., Jain, S., Kim, H. & Bar-Joseph, Z. Using neural networks for reducing the dimensions of single-cell RNA-Seq data. Nucleic Acids Res. 45, e156–e156 (2017).

35. Ma, J. et al. Using deep learning to model the hierarchical structure and function of a cell. Nat. Methods 15, 290–298 (2018).

36. Bhalla, U. S. & Iyengar, R. Emergent Properties of Networks of Biological Signaling Pathways. Science 283, 381–387 (1999).

37. Sachs, K., Perez, O., Pe’er, D., Lauffenburger, D. A. & Nolan, G. P. Causal Protein-Signaling Networks Derived from Multiparameter Single-Cell Data. Science 308, 523–529 (2005).

38. Jordan, J. D., Landau, E. M. & Iyengar, R. Signaling Networks: The Origins of Cellular Multitasking. Cell 103, 193–200 (2000).

39. Barabási, A.-L. & Oltvai, Z. N. Network biology: understanding the cell’s functional organization. Nat. Rev. Genet. 5, 101–113 (2004).

40. Srivastava, N., Hinton, G., Krizhevsky, A., Sutskever, I. & Salakhutdinov, R. Dropout: A Simple Way to Prevent Neural Networks from Overfitting. J Mach Learn Res 15, 1929–1958 (2014).

41. Datlinger, P. et al. Pooled CRISPR screening with single-cell transcriptome readout. Nat. Methods 14, 297–301 (2017).

42. Regev, A. et al. Science forum: the human cell atlas. Elife 6, e27041 (2017).

43. Yu, B. Stability. Bernoulli 19, 1484–1500 (2013).

44. Murdoch, W. J., Singh, C., Kumbier, K., Abbasi-Asl, R. & Yu, B. Interpretable machine learning: definitions, methods, and applications. ArXiv190104592 Cs Stat (2019).

45. Gillis, J., Ballouz, S. & Pavlidis, P. Bias tradeoffs in the creation and analysis of protein–protein interaction networks. J. Proteomics 100, 44–54 (2014).

46. Salvador, J. M. et al. Alternative p38 activation pathway mediated by T cell receptor-proximal tyrosine kinases. Nat. Immunol. 6, 390–395 (2005).

47. Falvo, J. V. et al. A stimulus-specific role for CREB-binding protein (CBP) in T cell receptor-activated tumor necrosis factor α gene expression. Proc. Natl. Acad. Sci. 97, 3925–3929 (2000).

48. Kim, H.-P. & Leonard, W. J. CREB/ATF-dependent T cell receptor–induced FoxP3 gene expression: a role for DNA methylation. J. Exp. Med. 204, 1543–1551 (2007).

49. Durant, L. et al. Diverse Targets of the Transcription Factor STAT3 Contribute to T Cell Pathogenicity and Homeostasis. Immunity 32, 605–615 (2010).

50. Thierfelder, W. E. et al. Requirement for Stat4 in interleukin-12-mediated responses of natural killer and T cells. Nature 382, 171–174 (1996).

51. Ellmeier, W. & Seiser, C. Histone deacetylase function in CD4 + T cells. Nat. Rev. Immunol. 18, 617 (2018).

52. Barndt, R. J., Dai, M. & Zhuang, Y. Functions of E2A-HEB Heterodimers in T-Cell Development Revealed by a Dominant Negative Mutation of HEB. Mol. Cell. Biol. 20, 6677–6685 (2000).

53. Woolf, E. et al. Runx3 and Runx1 are required for CD8 T cell development during thymopoiesis. Proc. Natl. Acad. Sci. 100, 7731–7736 (2003).

54. Ono, M. et al. Foxp3 controls regulatory T-cell function by interacting with AML1/Runx1. Nature 446, 685–689 (2007).

55. Herranz, D. et al. A NOTCH1-driven *MYC* enhancer promotes T cell development, transformation and acute lymphoblastic leukemia. Nat. Med. 20, 1130–1137 (2014).

56. Raaphorst, F. M. et al. Distinct BMI-1 and EZH2 Expression Patterns in Thymocytes and Mature T Cells Suggest a Role for Polycomb Genes in Human T Cell Differentiation. J. Immunol. 166, 5925–5934 (2001).

57. Gray, S. M., Amezquita, R. A., Guan, T., Kleinstein, S. H. & Kaech, S. M. Polycomb Repressive Complex 2-Mediated Chromatin Repression Guides Effector CD8+ T Cell Terminal Differentiation and Loss of Multipotency. Immunity 46, 596–608 (2017).

58. Murray, P. J. The JAK-STAT Signaling Pathway: Input and Output Integration. J. Immunol. 178, 2623–2629 (2007).

59. Stark, G. R. & Darnell, J. E. The JAK-STAT Pathway at Twenty. Immunity 36, 503–514 (2012).

60. Poli, V. The Role of C/EBP Isoforms in the Control of Inflammatory and Native Immunity Functions. J. Biol. Chem. 273, 29279–29282 (1998).

61. Gerritsen, M. E. et al. CREB-binding protein/p300 are transcriptional coactivators of p65. Proc. Natl. Acad. Sci. 94, 2927–2932 (1997).

62. Lécuyer, E. et al. The SCL complex regulates c-kit expression in hematopoietic cells through functional interaction with Sp1. Blood 100, 2430–2440 (2002).

63. Beurel, E., Michalek, S. M. & Jope, R. S. Innate and adaptive immune responses regulated by glycogen synthase kinase-3 (GSK3). Trends Immunol. 31, 24–31 (2010).

64. Macian, F. NFAT proteins: key regulators of T-cell development and function. Nat. Rev. Immunol. 5, 472–484 (2005).

65. Zhang, P. et al. Negative cross-talk between hematopoietic regulators: GATA proteins repress PU.1. Proc. Natl. Acad. Sci. 96, 8705–8710 (1999).

66. Porcher, C. et al. The T Cell Leukemia Oncoprotein SCL/tal-1 Is Essential for Development of All Hematopoietic Lineages. Cell 86, 47–57 (1996).

67. Nguyen, H. Q., Hoffman-Liebermann, B. & Liebermann, D. A. The zinc finger transcription factor Egr-1 is essential for and restricts differentiation along the macrophage lineage. Cell 72, 197–209 (1993).

68. Kong, K. Y. et al. Expression of Scl in mesoderm rescues hematopoiesis in the absence of Oct-4. Blood 114, 60–63 (2009).

69. Sarkar, A. & Hochedlinger, K. The Sox Family of Transcription Factors: Versatile Regulators of Stem and Progenitor Cell Fate. Cell Stem Cell 12, 15–30 (2013).

70. Liao, X. et al. Krüppel-like factor 4 regulates macrophage polarization. J. Clin. Invest. 121, 2736–2749 (2011).

71. Wurster, A. L., Tanaka, T. & Grusby, M. J. The biology of Stat4 and Stat6. Oncogene 19, 2577–2584 (2000).

72. Long, F. Building strong bones: molecular regulation of the osteoblast lineage. Nat. Rev. Mol. Cell Biol. 13, 27–38 (2012).

73. Vaillant, F., Blyth, K., Andrew, L., Neil, J. C. & Cameron, E. R. Enforced Expression of Runx2 Perturbs T Cell Development at a Stage Coincident with β-Selection. J. Immunol. 169, 2866–2874 (2002).

74. Schutten, E. A. et al. The role of Runx2 in CD8+ T cell memory during acute LCMV Armstrong infection. J. Immunol. 198, 78.8–78.8 (2017).

75. Taniuchi, I. et al. Differential Requirements for Runx Proteins in CD4 Repression and Epigenetic Silencing during T Lymphocyte Development. Cell 111, 621–633 (2002).

76. Milner, J. J. et al. Runx3 programs CD8+ T cell residency in non-lymphoid tissues and tumours. Nature 552, 253–257 (2017).

77. Kalev-Zylinska, M. L. et al. Runx3 is required for hematopoietic development in zebrafish. Dev. Dyn. 228, 323–336 (2003).

78. Wang, J. C. Y., Doedens, M. & Dick, J. E. Primitive Human Hematopoietic Cells Are Enriched in Cord Blood Compared With Adult Bone Marrow or Mobilized Peripheral Blood as Measured by the Quantitative In Vivo SCID-Repopulating Cell Assay. Blood 89, 3919–3924 (1997).

79. Shrikumar, A., Greenside, P. & Kundaje, A. Learning important features through propagating activation differences. in Proceedings of the 34th International Conference on Machine Learning-Volume 70 3145–3153 (JMLR. org, 2017).

80. Zeiler, M. D. & Fergus, R. Visualizing and Understanding Convolutional Networks. in Computer Vision – ECCV2014 (eds. Fleet, D., Pajdla, T., Schiele, B. & Tuytelaars, T.) 818–833 (Springer International Publishing, 2014).

81. Yosinski, J., Clune, J., Nguyen, A., Fuchs, T. & Lipson, H. Understanding Neural Networks Through Deep Visualization. ArXiv150606579 Cs (2015).

82. Simonyan, K., Vedaldi, A. & Zisserman, A. Deep Inside Convolutional Networks: Visualising Image Classification Models and Saliency Maps. ArXiv13126034 Cs (2013).

83. Liu, F., Li, H., Ren, C., Bo, X. & Shu, W. PEDLA: predicting enhancers with a deep learning-based algorithmic framework. Sci. Rep. 6, 28517 (2016).

84. Stegle, O., Teichmann, S. A. & Marioni, J. C. Computational and analytical challenges in single-cell transcriptomics. Nat. Rev. Genet. 16, 133–145 (2015).

85. Stuart, T. & Satija, R. Integrative single-cell analysis. Nat. Rev. Genet. 20, 257 (2019).

86. Aldridge, B. B., Burke, J. M., Lauffenburger, D. A. & Sorger, P. K. Physicochemical modelling of cell signalling pathways. Nat. Cell Biol. 8, 1195 (2006).

87. Raue, A. et al. Lessons Learned from Quantitative Dynamical Modeling in Systems Biology. PLOS ONE 8, e74335 (2013).

88. Scarselli, F., Gori, M., Tsoi, A. C., Hagenbuchner, M. & Monfardini, G. The graph neural network model. IEEE Trans. Neural Netw. 20, 61–80 (2008).

89. Wu, Z. et al. A comprehensive survey on graph neural networks. ArXiv Prepr. ArXiv190100596 (2019).

90. Dutil, F., Cohen, J. P., Weiss, M., Derevyanko, G. & Bengio, Y. Towards gene expression convolutions using gene interaction graphs. ArXiv Prepr. ArXiv180606975 (2018).

91. Rouillard, A. D. et al. The harmonizome: a collection of processed datasets gathered to serve and mine knowledge about genes and proteins. Database 2016, (2016).

92. Han, H. et al. TRRUST v2: an expanded reference database of human and mouse transcriptional regulatory interactions. Nucleic Acids Res. 46, D380–D386 (2018).

93. Perfetto, L. et al. SIGNOR: a database of causal relationships between biological entities. Nucleic Acids Res. 44, D548–D554 (2016).

94. Bateman, A. et al. UniProt: the universal protein knowledgebase. Nucleic Acids Res. 45, D158–D169 (2017).

95. Robinson, D. G. & Storey, J. D. subSeq: Determining Appropriate Sequencing Depth Through Efficient Read Subsampling. Bioinformatics 30, 3424–3426 (2014).

96. Abadi, M. et al. Tensorflow: a system for large-scale machine learning. in OSDI 16, 265–283 (2016).

97. Goodfellow, I., Bengio, Y., Courville, A. & Bach, F. Deep Learning. (The MIT Press, 2016).

98. Kuleshov, M. V. et al. Enrichr: a comprehensive gene set enrichment analysis web server 2016 update. Nucleic Acids Res. 44, W90–W97 (2016).

