## Supplemental Material for "Knowledge-primed neural networks enable biologically interpretable deep learning on single-cell sequencing data"

### SUPPLEMENTAL FIGURES

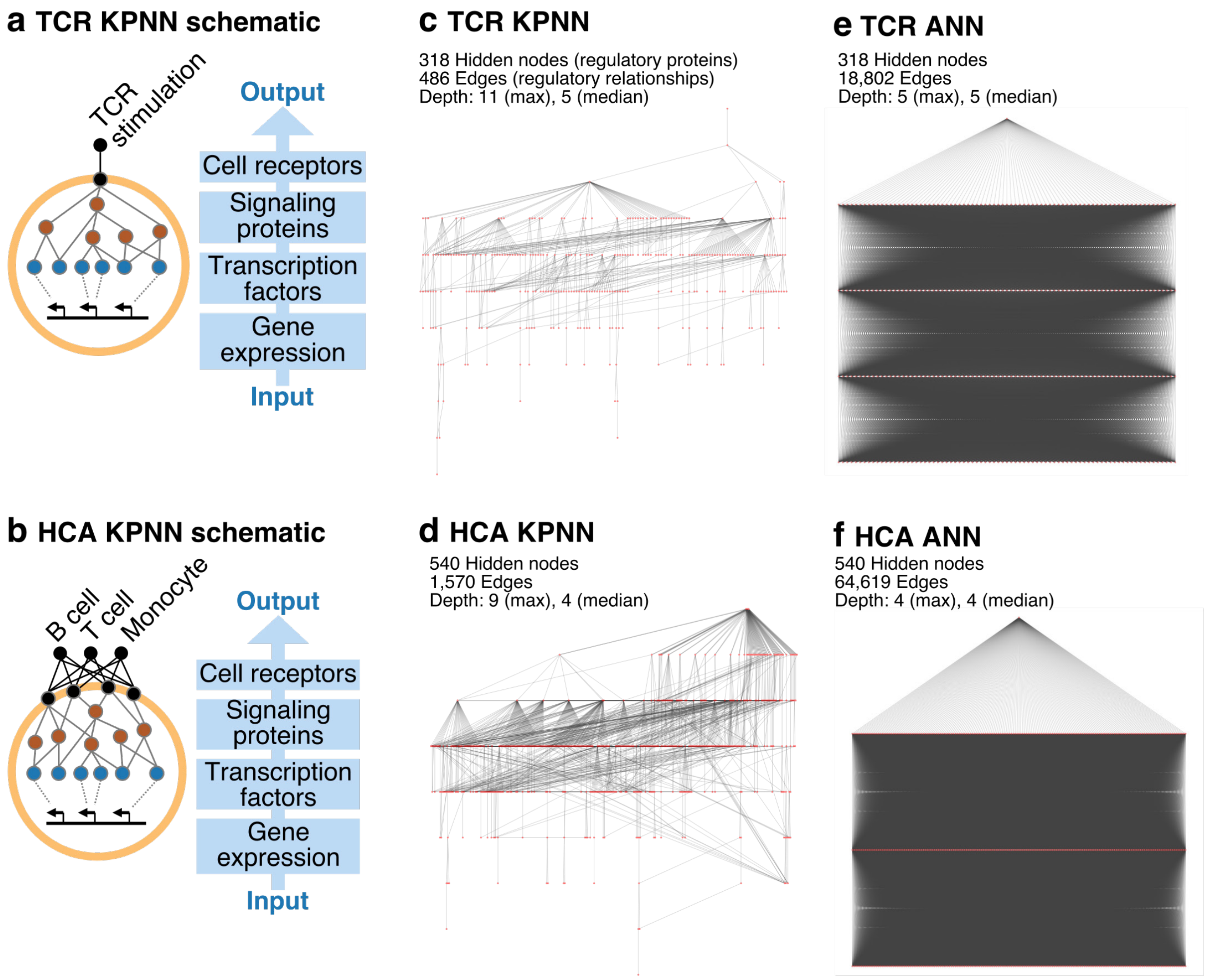


**Supplemental Figure 1**. Structure of KPNNs and their matched ANNs used for the TCR dataset **(a,c,e)** and the HCA dataset **(b,d,f)**. **(a,b)** Schematic outline of the two KPNNs. Based on public databases of signaling pathways and gene-regulatory networks, cell surface receptors are linked via signaling pathways to transcription factors, which are further linked to their target genes. Gene expression data are used as inputs to the learning algorithm. In the TCR KPNN **(a)**, the T cell receptor (TCR) is used as the output node to predict whether or not a T cell has been TCR stimulated. In the HCA KPNN **(b)**, cell type is predicted from all annotated cell surface receptors based on Human Cell Atlas (HCA) data for B cells, T cells, and monocytes. **(c)** Structure of the TCR KPNN, which predicts TCR stimulation from single-cell gene expression profiles. **(d)** Structure of the HCA KPNN, which predicts cell type from gene expression in a Human Cell Atlas dataset of immune cells. **(e-f)** ANNs with the same number of nodes and the same median depth as the TCR KPNN **(e)** and the HCA KPNN **(f)**. In all networks **(c-f)**, only output nodes and hidden nodes are shown (input nodes were removed for visualization). Network depth is calculated as the distance from the output node(s) to input nodes.

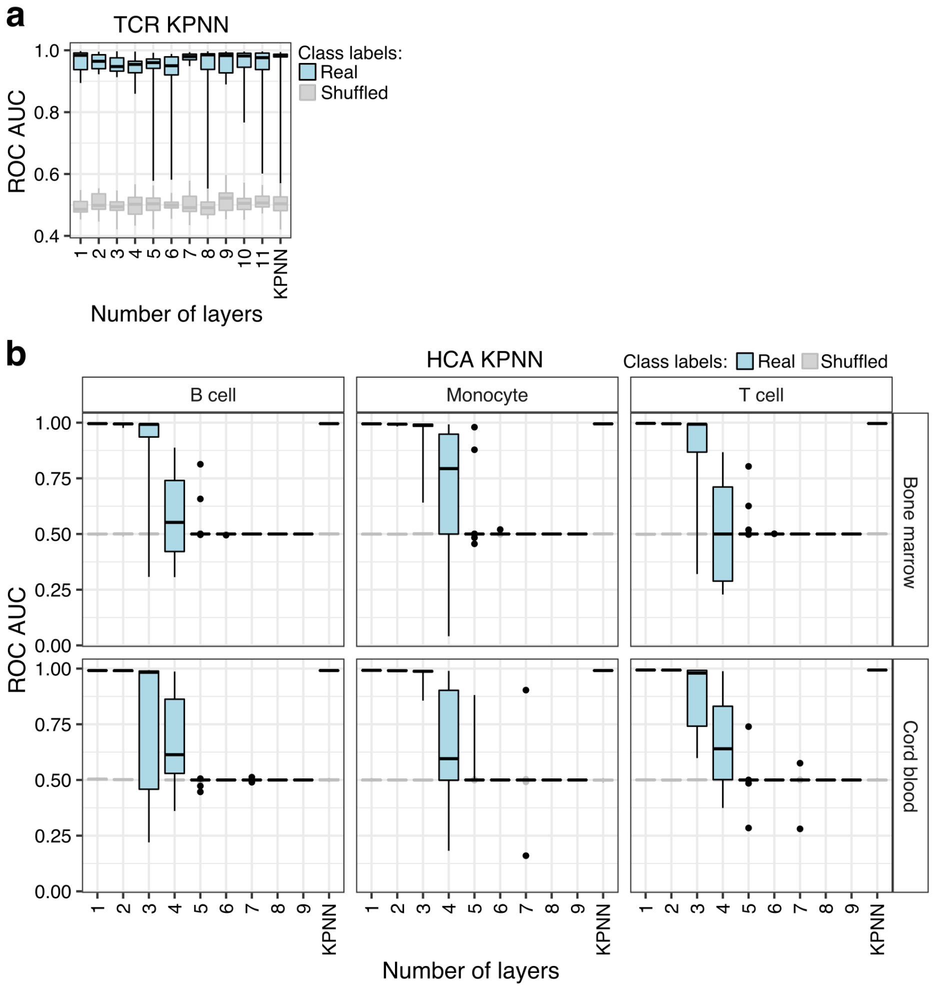


**Supplemental Figure 2**. Test set prediction performance of ANNs and KPNNs. KPNNs are compared to ANNs with the same number of nodes but different numbers of hidden layers. **(a)** ROC AUC values for the prediction of cell stimulation in the TCR KPNN. (**b**) ROC AUC for the prediction of cell type in the HCA KPNN. Values are shown separately for the HCA KPNN trained on data from bone marrow and from cord blood.

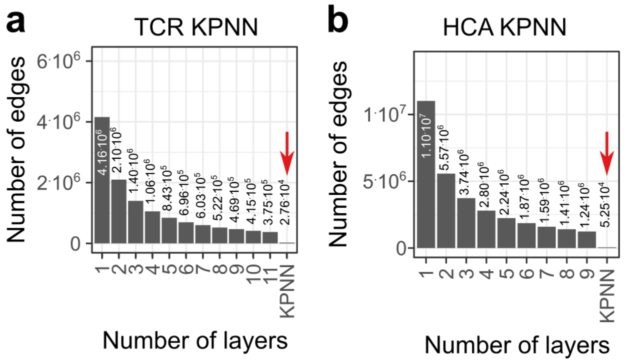


**Supplemental Figure 3**. Comparison of the number of edges between KPNNs and matched ANNs. **(a)** Number of edges for the TCR KPNN and for matched ANNs with the same number of nodes but different numbers of hidden layers. **(b)** Number of edges for the HCA KPNN and for matched ANNs with the same number of nodes but different numbers of hidden layers.

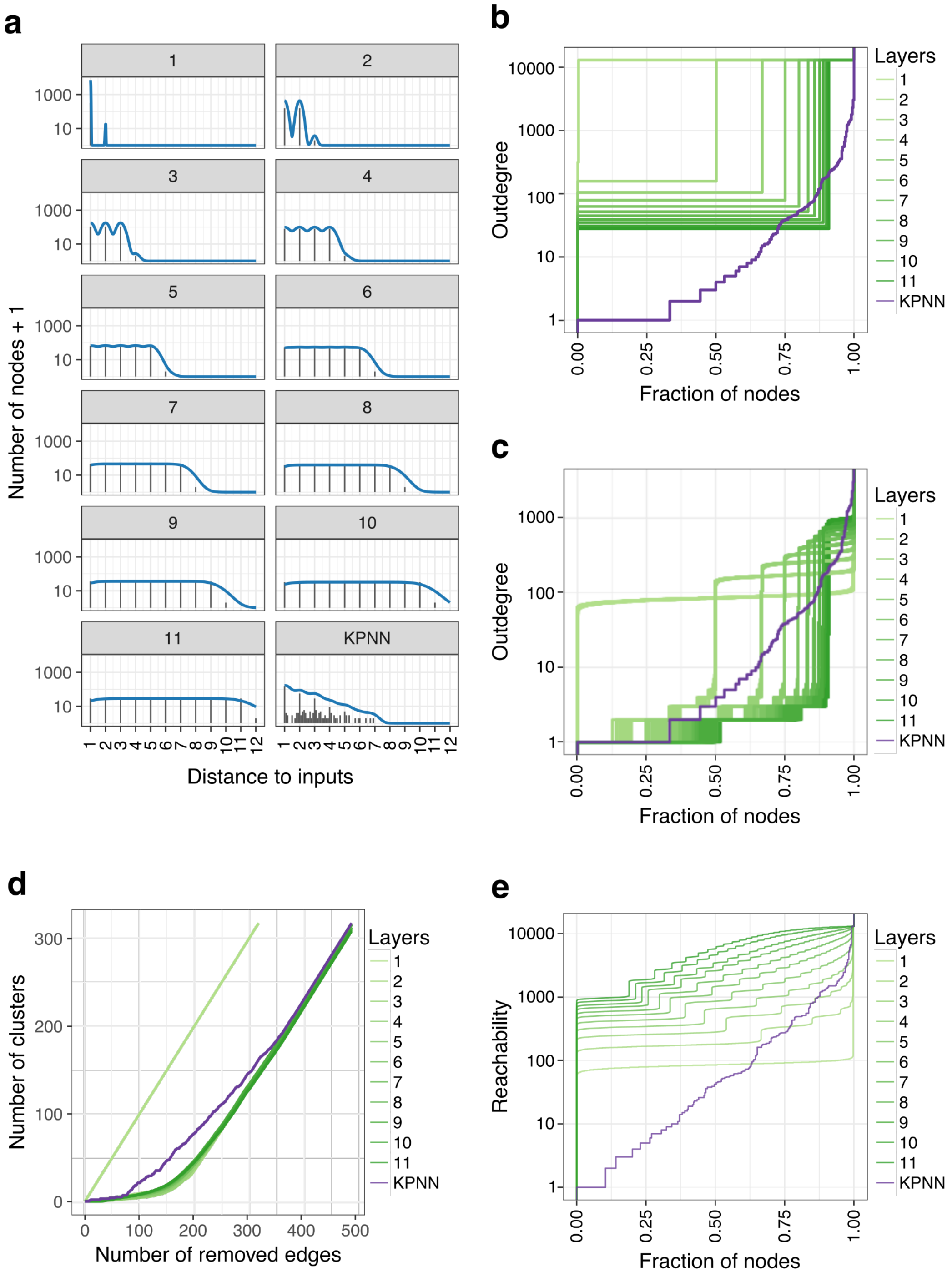


**Supplemental Figure 4**. Comparison of network structure of the untrained TCR KPNN and matched ANNs. The TCR KPNN is compared to fully connected ANNs (fANNs) with the same number of nodes as the KPNN, and with sparse ANNs (sANNs, n = 50 networks) where edges were randomly removed to resemble the KPNN. **(a)** Distribution of network distances (average distance of each hidden node to all input nodes) in the KPNN and in the fANNs. **(b)** Cumulative distribution of outdegree (number of downstream neighbors of each node) of hidden nodes in KPNN and in the fANNs. **(c)** Same as (b) for the KPNN and the sANNs. **(d)** Network sensitivity to targeted edge removal in the KPNN and the sANNs. Edges were removed based on their betweenness centrality values, and the number of disconnected clusters is plotted after removal of each edge. **(e)** Cumulative distribution of reachability of hidden nodes in the KPNN and in the sANNs. Reachability values measure the number of input nodes that a hidden node can access directly or indirectly (i.e., the number of inputs that are available to the hidden node).

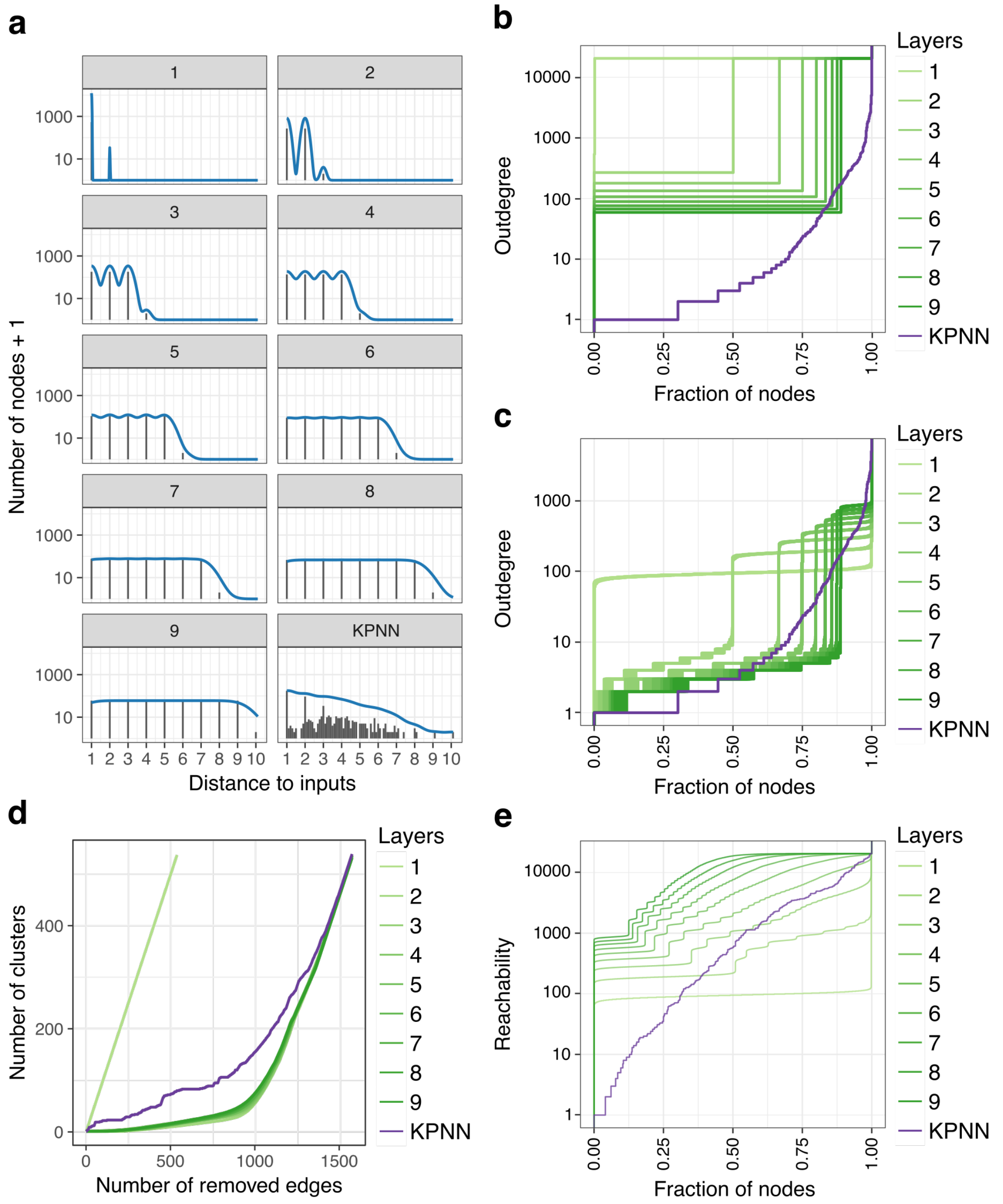


**Supplemental Figure 5**. Comparison of network structure of the untrained HCA KPNN and matched ANNs. The HCA KPNN is compared to fully connected ANNs (fANNs) with the same number of nodes as the KPNN, and with sparse ANNs (sANNs, n = 50 networks) where edges were randomly removed to resemble the KPNN. **(a)** Distribution of network distances (average distance of each hidden node to all input nodes) in the KPNN and in the fANNs. **(b)** Cumulative distribution of outdegree (number of downstream neighbors of each node) of hidden nodes in KPNN and in the fANNs. **(c)** Same as (b) for the KPNN and the sANNs. **(d)** Network sensitivity to targeted edge removal in the KPNN and the sANNs. Edges were removed based on their betweenness centrality values, and the number of disconnected clusters is plotted after removal of each edge. **(e)** Cumulative distribution of reachability of hidden nodes in the KPNN and in the sANNs. Reachability values measure the number of input nodes that a hidden node can access directly or indirectly (i.e., the number of inputs that are available to the hidden node).

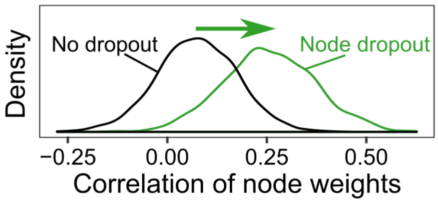


**Supplemental Figure 6**. Effect of dropout on node weight correlations. Density plots are shown for the correlation of node weights learned with and without hidden node dropout.

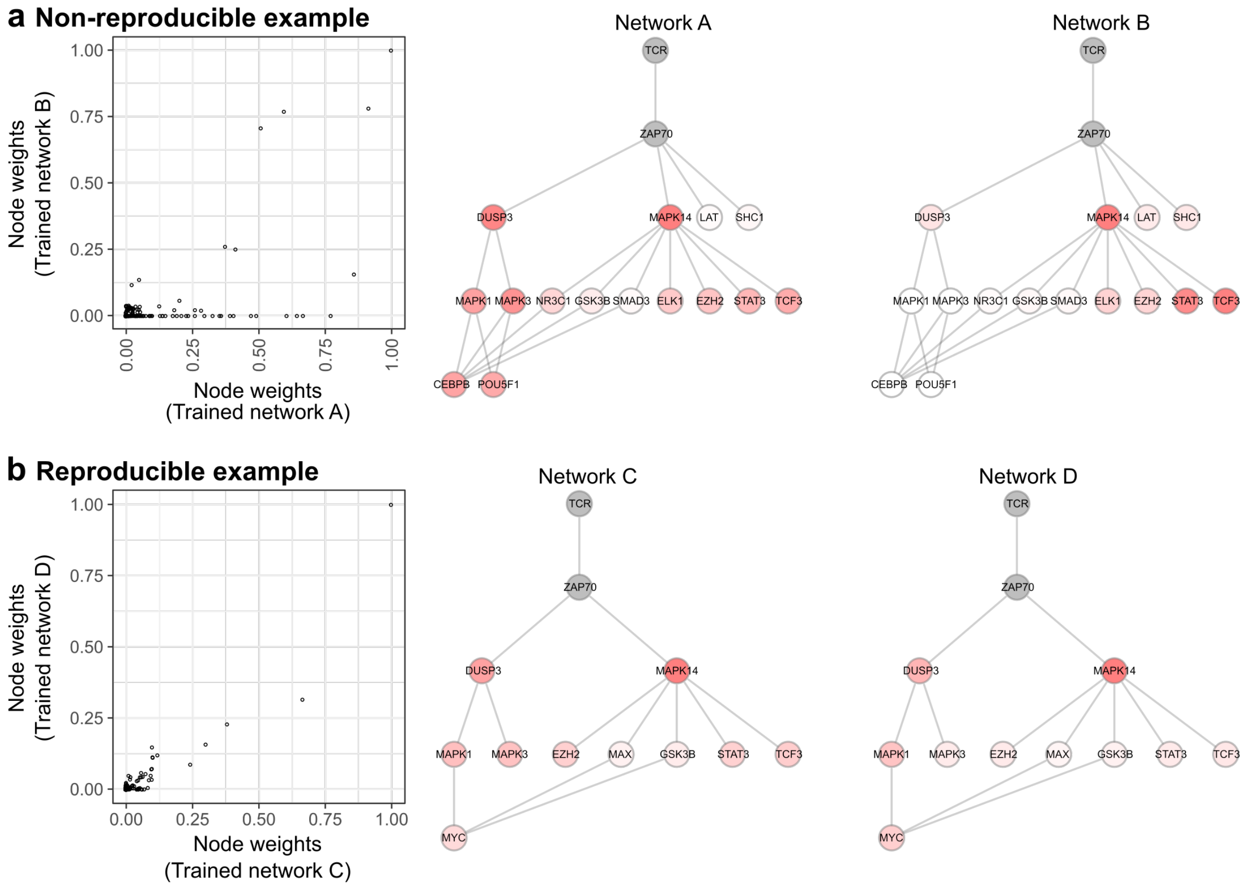


**Supplemental Figure 7**. Detrimental effect of low reproducibility on interpretability. **(a)** Example of low correlation between two network replicates learned without dropout. The two networks produce highly inconsistent results. This is exemplified by the node weights for DUSP3, which receives high importance in only one of the two replicate networks. **(b)** Example of high correlation between two replicates learned with dropout.

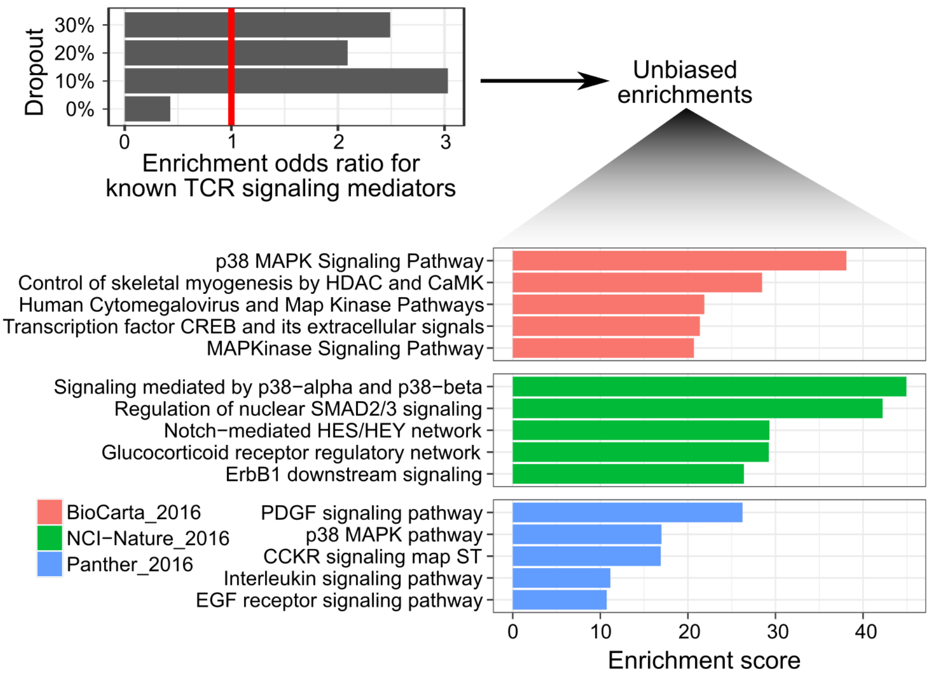


**Supplemental Figure 8**. Interpretation of node weights in the TCR KPNN. (**Top**) Enrichment of annotated TCR signaling mediators among significantly differential nodes at different dropout rates. (**Bottom**) Gene set enrichment analysis of differential nodes at a dropout rate of 10%. The five most enriched gene sets from each database are shown.

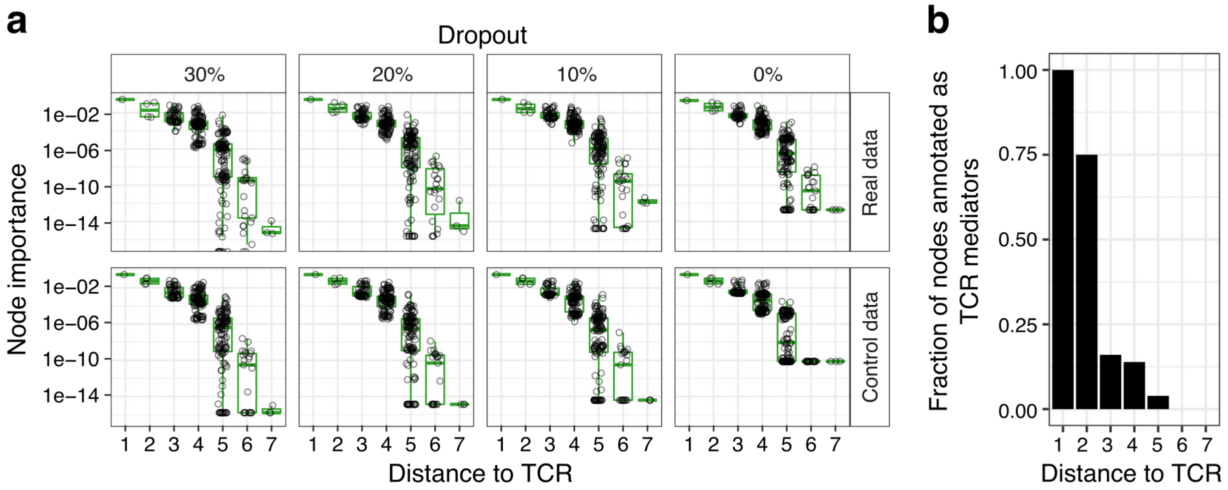


**Supplemental Figure 9**. Effect of the uneven connectivity of biological networks on node weights in the TCR KPNN. (**a**) Relationship between node weights and the distance to the output node for the TCR KPNN. Hidden nodes closer to the output node have a greater effect on network output and therefore receive larger node weights not only on the actual data, but also on control inputs. (**b**) Nodes closest to the TCR in the KPNN are most strongly enriched for annotated TCR mediators.

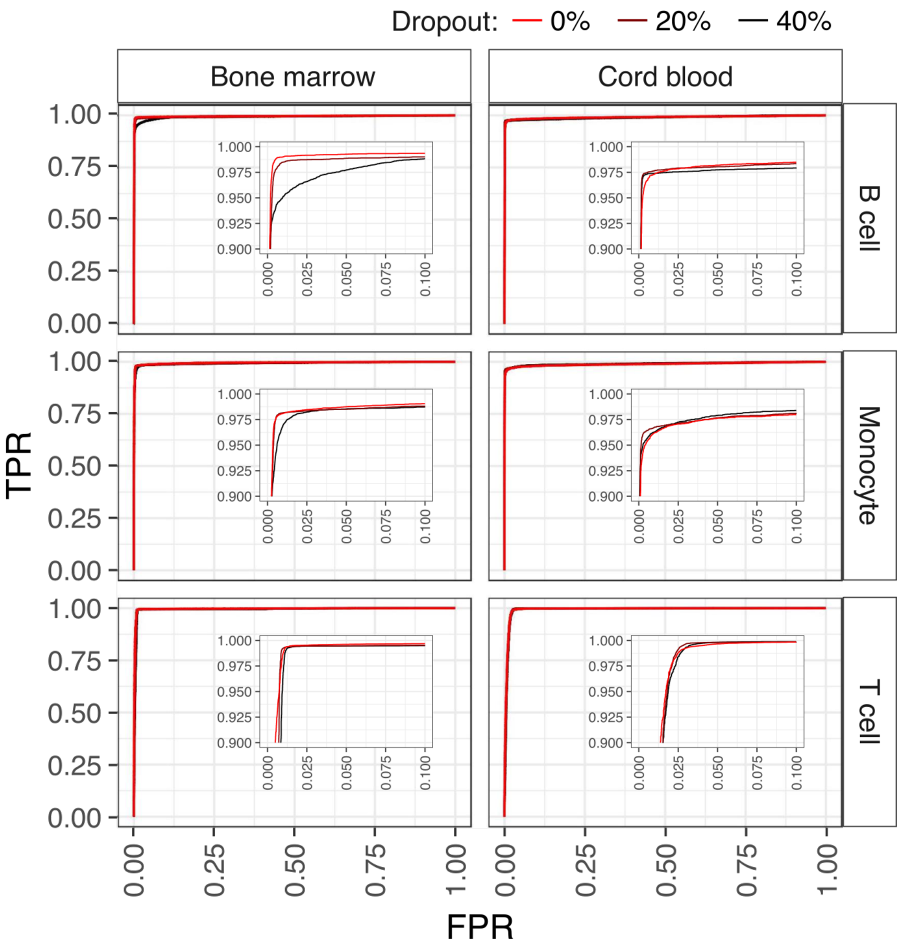


**Supplemental Figure 10**. Prediction accuracy of the HCA KPNN. ROC curves are shown for networks trained with different levels of dropout applied to hidden nodes as well as input nodes. Curves for the HCA KPNN with ROC AUC closest to the mean at each dropout rate are shown. The inset shows a zoomed-in view.

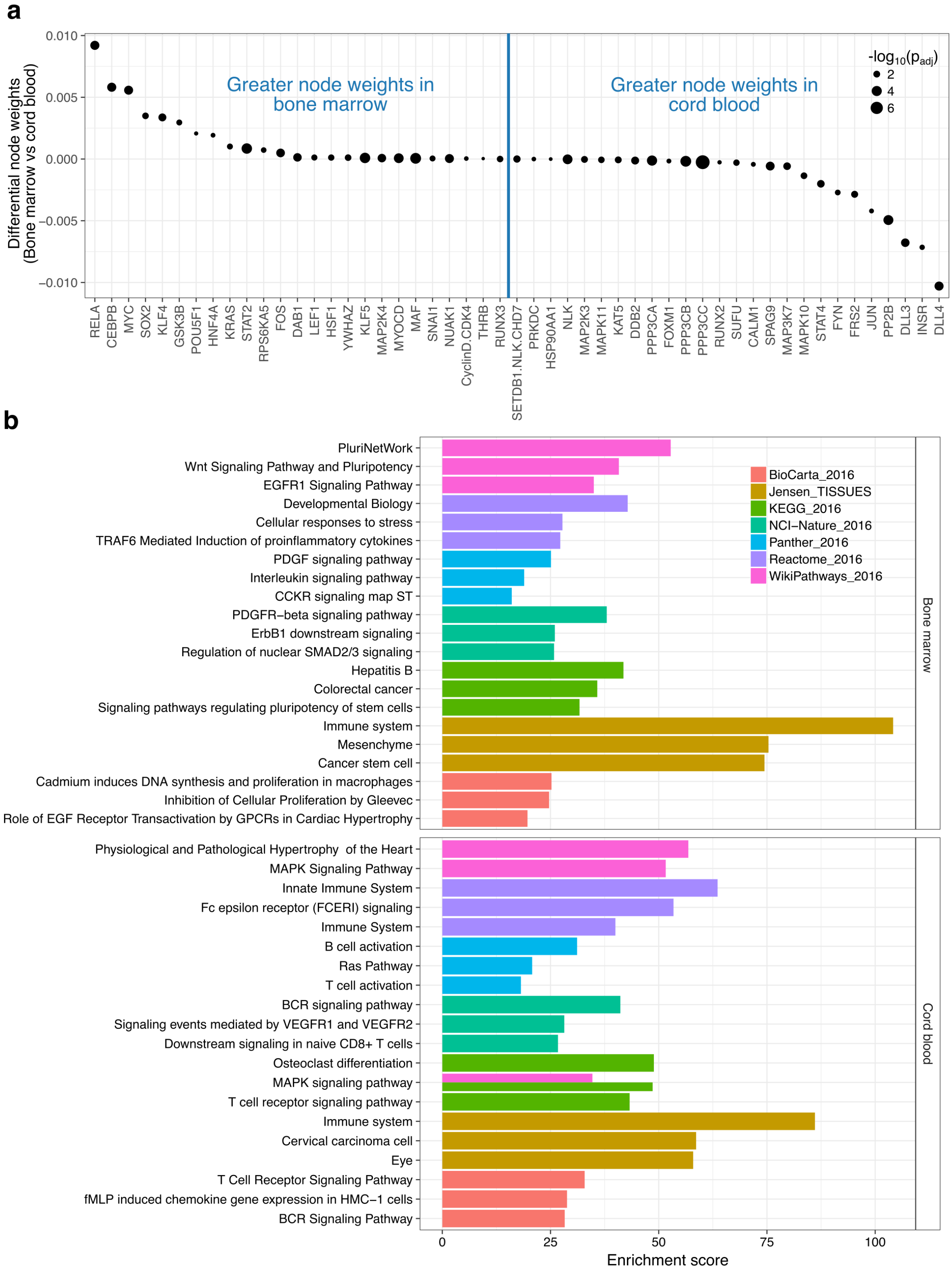


**Supplemental Figure 11**. Regulatory processes that distinguish immune cells in the bone marrow and cord blood. (**a**) Differential node weights between the two sources of analyzed immune cells, bone marrow and cord blood. Nodes with significantly different weights ($p_{adj} < 0.05$) are shown. (**b**) Gene set enrichment analysis of source-specific nodes. The three most enriched gene sets from each database are shown for both sources.

### SUPPLEMENTAL TABLES

**Supplemental Table 1**. Differential node weights for the TCR KPNN trained on actual data and on control inputs.

**Supplemental Table 2**. Results of an Enrichr analysis for differential node weights from the TCR KPNN.

**Supplemental Table 3**. Differential node weights for the HCA KPNN trained on actual data and on control inputs.

**Supplemental Table 4**. Differential node weights for the HCA KPNN trained on actual data comparing bone marrow and cord blood.

**Supplemental Table 5**. Results of an Enrichr analysis for differential node weights from the HCA KPNN comparing bone marrow and cord blood.

### SUPPLEMENTAL VIDEOS

**Supplemental Video 1**. Effect of dropout on the robustness of learned weights in KPNNs. The video visualizes the learning process for the TCR KPNN, trained twice without dropout (standard learning method) (**top**) and twice with dropout (optimized learning method) (**bottom**). Edge color and thickness reflect edge weights, with thick red lines indicating high weights and thin grey lines indicating low weights. Edge weights were transformed to absolute values prior to plotting. The networks shown were selected by first correlating edge weights between all pairs of networks, and then selecting the two networks that are closest to the median of all pairwise correlations (R = 0.78 without dropout and R = 0.86 with dropout).
